# Revisiting bacterial division control with likelihood inference: limited identifiability of simple models

**DOI:** 10.64898/2026.01.16.700002

**Authors:** Ron Teichner, Ron Meir, Naama Brenner

## Abstract

Cell-division control in bacteria has been studied for many years, but gaps in understanding its logic still remain. Simple candidate models of cell division control have been studied, but are assessed with heuristic analysis of single-cell data that does not provide a quantitative scale for comparison. We recast division control mechanism identification as a likelihood-based inference problem, defining an explicit metric for model comparison: among candidate mechanisms, the model with higher likelihood provides the better explanation of the data. We demonstrate that within a broad class of models, discrimination depends only on the conditional distribution of cell-cycle durations. Under mild independence assumptions, variability in growth rates and division asymmetry do not contribute to the likelihood-based comparison, effectively separating the problem of understanding growth from that of identifying division control. Applying the likelihood framework to simulations and experimentally measured long-term single-cell lineages of *Escherichia coli*, we find that candidate simple models are statistically identifiable in principle, but only weakly separated in practice. Sizer, adder, and related mechanisms all achieve comparable and generally high likelihoods, but no single one consistently outperforms others: their relative likelihood depends on the dataset and growth condition. In contrast, a flexible model trained directly on the data always achieves significantly higher likelihood, revealing structure not captured by any one of the simplified rules; surprisingly, it can be well fit by a low-dimensional variable combination. More broadly, this work establishes a general likelihood-based framework for identifying stochastic regulatory mechanisms and for comparing future mechanistic models on a quantitative scale.

## I. Introduction

Biological regulation is often inferred indirectly, through noisy observables shaped by multiple interacting sources of variability. A central challenge in systems biology is therefore not only to describe phenotypic regularities, but to identify which variables are actively controlled and how competing mechanistic hypotheses should be compared. Cell division provides a particularly important instance of this problem: Proliferating cells grow and divide over many generations while maintaining stable distributions of all properties despite substantial stochastic fluctuations, raising the question of which variables govern division and whether a control rule can be inferred from data.

Lineages of growing and dividing bacteria offer a natural setting for addressing this question. Recent advances in microfluidics and time-lapse microscopy have enabled long single-cell lineage measurements with sufficient temporal resolution to quantify birth size, division size, growth rate, and cycle duration across many consecutive generations [3], [18], [20], [22], [23], [25]–[27]. These data have motivated several simple models as candidate mechanisms for division control: in the *sizer* model, cells divide upon reaching a critical size; in the *adder* model, division is triggered after adding an approximately fixed size increment; and in the *log-foldchange* (LFC) model, division is associated with reaching a target multiplicative growth factor over the cell cycle [1], [9], [19]. Distinguishing among these mechanisms can advance our understanding of how cells achieve growth-and-division homeostasis, yet it remains unclear whether any of them fully captures the true biological control rule.

A longstanding difficulty is that mechanism identification has typically relied on low-dimensional statistical features, such as correlations between birth size and added size [1]. Such heuristics are intuitively appealing, but do not provide a fully defined metric for comparing competing mechanisms, and their interpretation can depend strongly on modeling assumptions. In particular, Luo et al. [12] showed that classification based on these correlations is valid only when the division threshold is fixed. If the threshold itself fluctuates dynamically, the same underlying mechanism can generate markedly different correlation patterns, depending on the timescale of these fluctuations. Thus, widely used heuristics can become unreliable precisely in the stochastic regime most relevant to living cells.

Here we address this problem by formulating division-control mechanism identification as a maximum-likelihood inference problem. Candidate mechanisms are compared on a common quantitative scale and ranked according to the relative likelihood they assign to the data. We find that, within a broad class of models, this comparison reduces to the relative likelihood of observed cell-cycle durations.

We apply the framework to both simulated and experimental data. On simulated datasets, it reliably distinguishes the ground-truth mechanism among competing candidate models, demonstrating that a division control mechanism is statistically identifiable in principle in the stochastic regime considered here; however, we find that in practice they are only weakly separated. Applying the framework to six *E. coli* datasets reveals that indeed previously suggested candidate models are only weakly distinguishable, and remain informative approximations even when not ranked highest. Moreover, no single one of these mechanisms is universally most likely: their relative ranking depends on growth condition and dataset. In contrast, a flexible data-driven model trained directly on experimentally measured lineages, achieves the highest likelihood in all cases examined. Interestingly, this model can be well approximated by a low-dimensional combination of variables, different from the previously suggested simple control rules, suggesting that additional structure remains to be uncovered beyond these rules.

## II. From heuristic classifications to a likelihood-based framework

Typical single-cell growth and division trajectories of bacteria measured in microfluidic mother-machine devices exhibit approximately exponential accumulations separated by abrupt, noisy symmetric divisions. In such a lineage trajectory, only the mother cell is followed over time; for each cycle *k* we observe the set of measurements

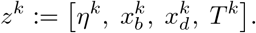

At the start of cell cycle *k* birth time 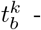 the cell size is 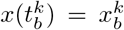 and at division time,is the division fraction. Within each cycle, therefore, a good approximation to the trajectory is

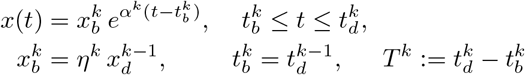

where *T*^*k*^ is the cycle duration. The relations between the cycle parameters are thus

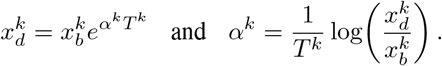

Exponential growth law within a cycle, with a variable rate *α*^*k*^, is a parsimonious and empirically supported approximation [18], [20], [25].

Figure 1 (top) illustrates simulations of such trajectories, where *α*^*k*^ is drawn each cycle from some given distribution. Simple models of cell division control describe the division event as occuring when a division indicator crosses a fluctuating threshold *u*(*t*) - brown line. In the figure, this indicator is cell size (left column), and added cell size (right column), the sizer and adder models respectively [1], [9].

**Fig. 1.**
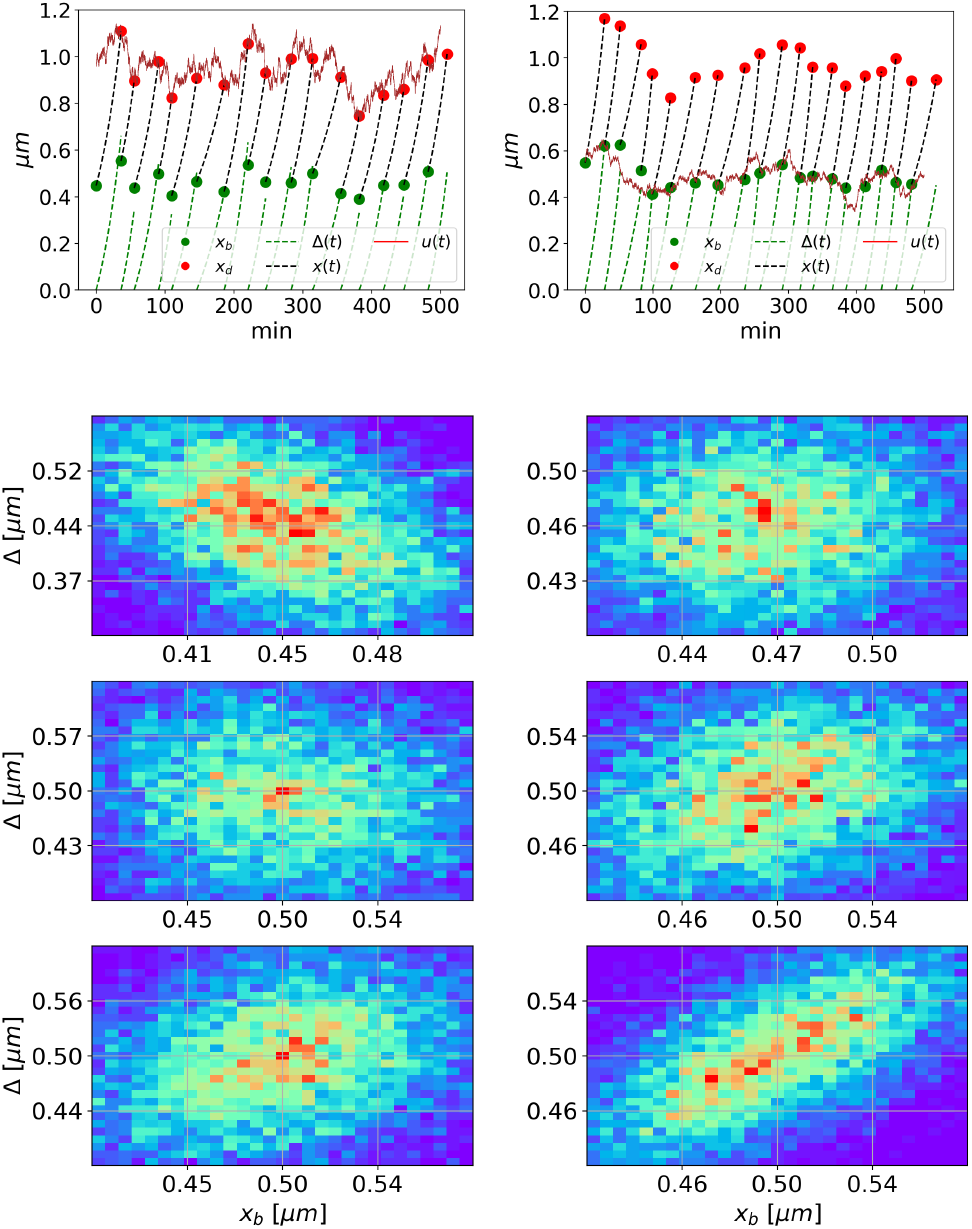
**Top**: Two simulated lineages generated by stochastic-threshold models. After abrupt division, cell size starts from size *x*_*b*_ (green circle) and grows exponentially (dashed black line) until the next division (red circle); added size Δ(*t*) is depicted as a dashed green line. Note that each division gives rise to two daughter cells, whereas only one daughter is followed in the lineage plot. The threshold *u*(*t*) (brown line) follows an Ornstein-Uhlenbeck process with correlation time *τu* = 40 min, close to the average cycle time. Parameters (*μu, σu*) of the threshold: (1, 0.1) (left), (0.5, 0.05) (right; see Appendix A-C for formulas). **Bottom**: Correlation plots of added size (Δ = *x*_*d*_ − *x*_*b*_) versus birth size (*x*_*b*_), illustrating the dependence of the heuristic identification method on the threshold autocorrelation time *τ*^*u*^ (*τ*^*u*^ ≪ E[*T*] at top row, *τ*^*u*^ ≈ E[*T*] middle and *τ*^*u*^ ≫E[*T*] at bottom). Left column: sizer mechanism. Right column: adder mechanism. Only when *τ*^*u*^ = 2 min ≪ E[*T*] do these plots correctly identifies the division control model.

To identify which mechanism is at work in lineage data, a common approach is to examine the correlation between birth size *x*_*b*_ and added size Δ = *x*_*d*_ − *x*_*b*_. Data are classified as following a sizer mechanism when Δ is negatively correlated with *x*_*b*_, and as following an adder mechanism when the two are approximately uncorrelated [1]. However, such heuristics are valid only under restrictive assumptions [12]. Figure 1-bottom illustrates this limitation: In the top row, threshold correlation time *τ*_*u*_ is much smaller than average cycle time E[*T*], and the correlations display the expected behavior. As the threshold correlation time is increased in the bottom two rows, this behavior changes and the correlation plots no longer reliably distinguish between mechanisms. This ambiguity motivated us to revisit the problem of division control identification from a different approach.

One may wonder whether lineage data of the type measured in experiments contains enough information to distinguish reliably between mechanisms. In this context it is useful to consider an asymptotic regime in which the identification problem becomes intrinsically non-informative. In the limit of a constant threshold (*u*(*t*) = const) and perfectly symmetric division, lineage trajectories converge to repeated cycles of identical size and added-size. In that regime, all candidate division models generate the same observations, and therefore explain the data equally well. Equivalently, the likelihoods under the competing hypotheses coincide, and the underlying mechanism is not statistically identifiable from the observations alone.

This limiting case clarifies the role of stochastic variability. Dynamic threshold fluctuations, growth-rate variability and division asymmetry act as perturbations away from the perfectly degenerate regime, and it is these perturbations that create the statistical signal needed to distinguish between mechanisms. In particular, *a-priori* neglect of fluctuations in any of the variables can strongly affect the results (as for example assuming a fixed threshold or negligibly small growth rate variability).

We will consider a general class of probabilistic models with stochastic fluctuations in all main variables. Randomness in *α*^*k*^ accounts for cell-to-cell variability through the conditional probability density

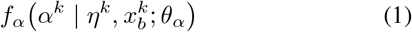

parameterized by *θ*_*α*_. For example, growth rates drawn independently from a Gamma distribution, *α*^*k*^ ∼ Γ(*θ*_*α*,1_, *θ*_*α*,2_), consistent with experimental observations [3], [16], is a special case used in our simulations. Cell division is approximately symmetric for many bacteria, and we model it as a Gaussian distribution around 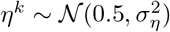

We have specified the growth and division model without reference to the division-control mechanism that we wish to identify. Such a mechanism, denoted by *γ*, comprises of the indicator that needs to cross the threshold for division to occur, and the statistical properties of the threshold itself. Each mechanism, with parameters *θ*_*γ*_, will result in a conditional distribution of cell cycle durations

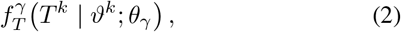

where we collect the conditioning observations in a single variable,

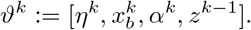

The key structural assumption we make is that the parameters 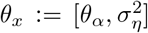 governing the growth law and asymmetry at division, are independent of the parameters that characterize the division control mechanism, *θ*_*γ*_, thus separating the model into two decoupled parts.

Our *inferential goal* is to determine the division indicator *γ* from the observations *z*^0:*N*−1^, while treating the parameters *θ*_*γ*_ and *θ*_*x*_ as unknown; two competing mechanisms will be compared at the best-fit parameter values of each one. To that end, we define the likelihood under each hypothesis *γ* as

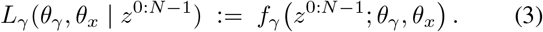

To compare two candidate mechanisms *γ* and *δ*, we should estimate the generalized likelihood ratio for observing the data

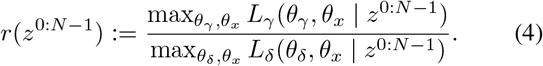

If *r* > 1, mechanism *γ* is favored over *δ*; if *r* < 1, mechanism *δ* is favored. When *r≈* 1, the data do not clearly discriminate between the two mechanisms. Because of the decomposition of the problem into two independent parts - growth/division model, division control mechanism - as described above, the generalized likelihood ratio (4) reduces to a quantity that only depends on the conditional distributions of cycle durations, while the parameters *θ*_*x*_ cancel from the ratio and do not contribute:

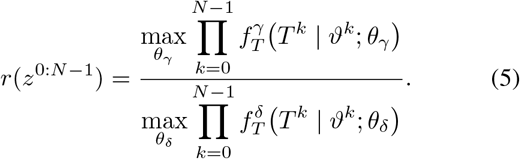

This result is formulated here in a simplified version, and proven for a more generalized class, that includes also history dependence, in Appendix Section A. It considerably simplifies the likelihood ratio, and renders the conditioned cycle time distribution 2 a central object in our analysis. For the LFC model, this distribution can be formulated in closed form, while for the adder and sizer models we approximate it by the generalized Gamma family, which consistently provides the best fit (see Materials and Methods Section IV-A).

The corresponding log-likelihood of division-control mechanism *γ* is defined for an individual observation *T*^*k*^,

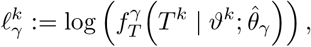

with 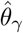 the best estimated parameters. Averaged over *N* cycles we define the log-likelihood of mechanism 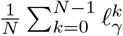 Both will be analyzed in the following sections.

For reference, we also fit an unconditional distribution directly to the observed cycle durations,

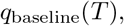

and define its per-cycle log-likelihood as 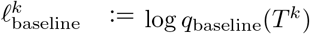 By construction, this baseline represents the marginal distribution of cycle durations. It therefore provides a benchmark, allowing to assess whether conditioning on the cell state provides an improvement over modeling only the marginal distribution of cycle duration. We next apply these comparisons to simulated and experimental data.

### A. Mechanism comparison on simulation data

We first ask whether the maximum-likelihood estimator 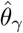recovers the ground-truth parameters *θ*_*γ*_ in simulations where the division mechanism is known. To answer that, we generated multiple simulations under the adder division mechanism, with parameters chosen in the biologically relevant regime (see Appendix Section B-5 and Proposition B.1). Figure 2 (top) shows the values of estimated parameters of the threshold process timescale (left) and standard deviation (right) - as a function of the ground truth values. The estimates are concentrated near the true parameters (up to an expected systematic bias in the standard deviation), indicating that the optimization procedure locates the correct region of parameter space. Similar results for other ground-truth division control mechanisms are shown in Appendix F-A.

**Fig. 2.**
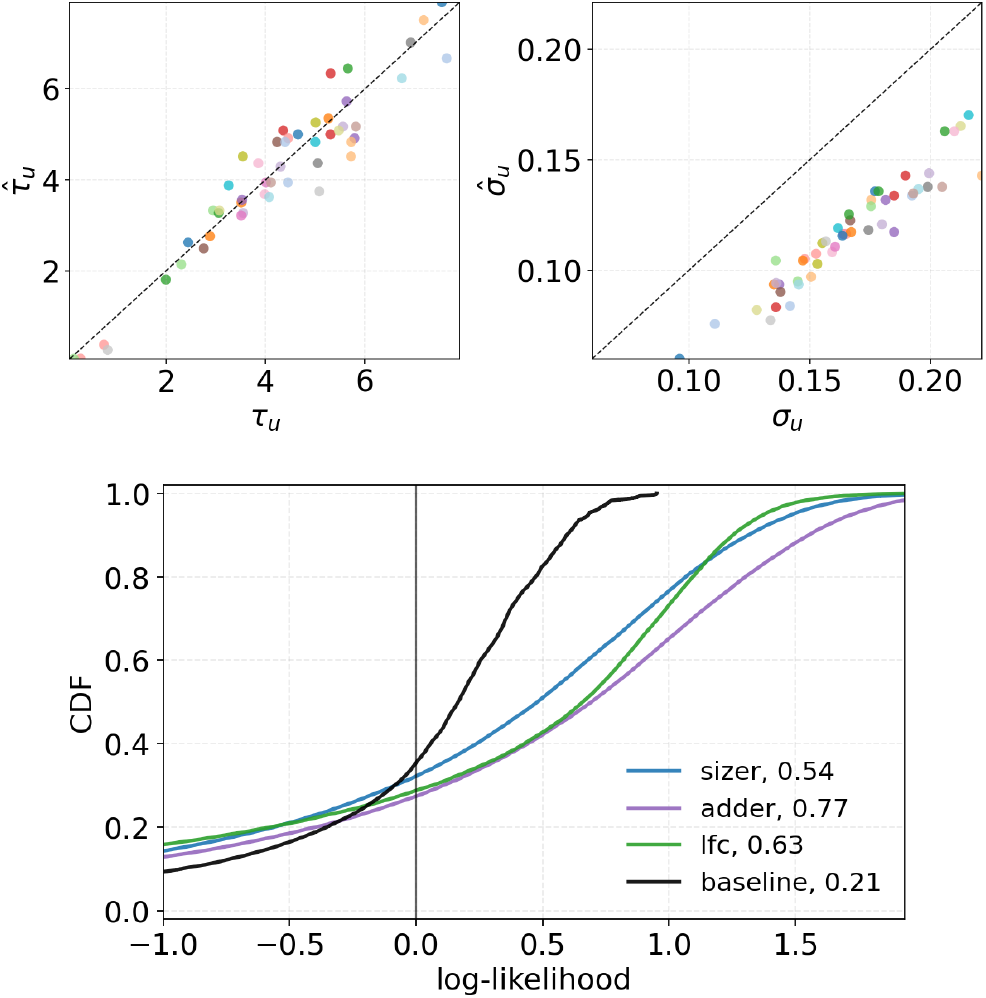
Applying likelihood inference to adder simulations. Each dataset includes 50 simulated lineages of 25 cycles, resembling experimental datasets, simulated with the adder control mechanism. **(Top)** Maximum-likelihood parameter estimates as a function of true simulation values: threshold correlation timescale *τ*^*u*^ (left), and threshold standard deviation *σ*^*u*^ (right). The average of the threshold is normalized to one (see Materials and Methods Section B-5). A systematic bias is visible in *σ*^*u*^, consistent with the known finite-sample bias identified by Hurwicz [7]. **(Bottom)** Cumulative distributions of per-cycle log-likelihood for competing candidate division-control mechanisms, each evaluated at its corresponding maximum-likelihood parameters. The legend presents the averaged log-likelihoods, 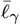, demonstrating that the adder mechanism achieves the highest average log-likelihood. The per-cycle log-likelihood values are highest for the adder mechanism across almost the entire quantile range, indicating that the adder’s advantage is broadly distributed rather than resulting from a small subset of observations

The second question is whether the ground truth mechanism receives higher likelihood than the alternative candidates.

Using the additive decomposition of the log-likelihood, we compare the empirical cumulative distribution functions (CDFs) of the per-cycle log-likelihoods 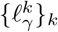 in Figure 2 (bottom). The per-cell log-likelihood of the true adder mechanism (purple line) dominates that of the competing mechanisms (other colors) across the entire range of quantiles. In other words, the adder mechanism has an advantage which is not dominated by a small number of extreme cells, but is distributed broadly across the observations. The average likelihood 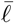 is the highest for the true adder mechanism [17];

averages are displayed in the legend. Importantly, however, competing candidate models can come quite close to the performance of the adder, exhibiting a relatively small likelihood gap. This reflects the fact that an incorrect mechanism may still provide a quantitatively good account of the data. Additional simulation results are presented in Appendix Section F-A.

Taken together, the simulation results show that the correct division-control mechanism is, in principle, statistically identifiable in the relevant stochastic regime the ground-truth is consistently assigned the highest likelihood, both at the level of total dataset likelihood and at the level of the per-cycle likelihood distribution. It calibrates our expectation with respect to the likelihood gap between competing mechanisms; this will be important when we next examine experimental data, for which the ground-truth mechanism is unknown.

### B. Mechanism comparison on experimental data

We next apply the likelihood-based framework to six long-term single-cell lineage datasets of *Escherichia coli*, obtained in microfluidic (mother machine) devices under different environmental conditions, spanning various temperatures, medium compositions, and strain backgrounds.

Our first observation is that no single division control mechanism universally best explains all experiments. Each dataset was fitted separately under each candidate mechanism, and the optimization was repeated 20 times to assess the stability of the resulting likelihood values. Fig. 3 (top) summarizes the average likelihoods with blue, purple, and green bars denoting the sizer, adder, and LFC models, respectively, and with error bars showing the standard deviation across runs. Importantly, the preferred mechanism depends on details even among experiments on the same strain performed in the same lab. For example, in the Tanouchi datasets [22], the two higher-temperature conditions are best explained by the sizer model, whereas the lowest-temperature condition favors LFC. Similarly, the Sassi datasets [16] slightly favor a sizer in rich medium but an adder in poor medium.

**Fig. 3.**
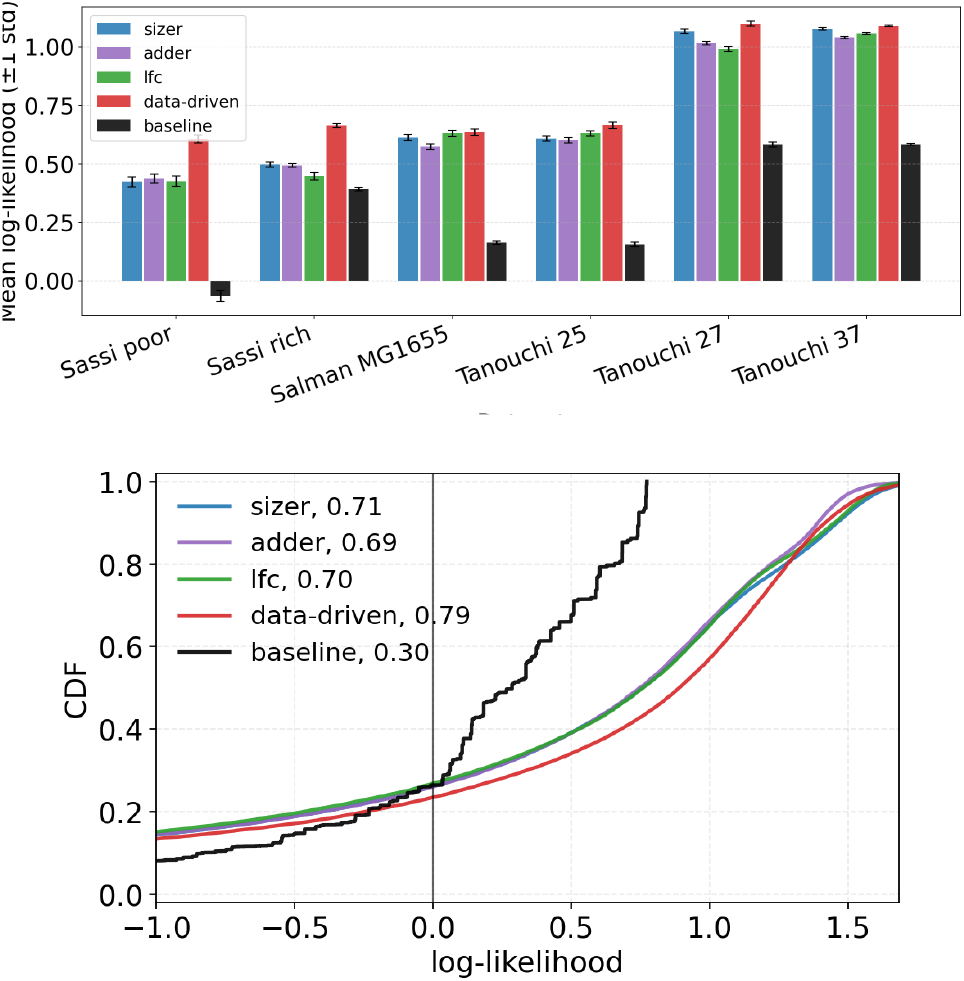
Applying likelihood inference to *E. coli* mother machine data. Six datasets were analyzed: ProVenus strains at 30^*°*^C under either poor (’Sassi poor’, supplemented M9) or rich (’Sassi rich’, MOPS EZ) nutrient conditions [16]; MG1655 strain at 30^*°*^C in LB medium (Salman MG1655’, [19]); MC4100 strain in LB medium at 25^*°*^C (’Tanouchi 25’) and similarly at 27^*°*^C and 37^*°*^C [21], [22]). **Top**: Log-likelihoods obtained by fitting each dataset separately under each candidate model. Bar height is the mean over 20 independent optimization runs; error bars denote standard deviation across runs. Numerical results are reported in Appendix Table F.2, and the corresponding fitted parameters - in Appendix Table F.1. **Bottom**: Distributions of per-cycle log-likelihoods under the competing models, evaluated for all data. Parameters are fitted separately for each data-set and each mechanism, and the distribution shows all likelihoods pooled together. reported in Appendix Table F.1. We note that the reported likelihoods were evaluated on randomly selected held-out cycles that were not used for model training, and hence reflect the true out-of-sample performance.

It is informative to compare these results with previous heuristic classifications of the same data. Tanouchi *et al*. [22] analyzed the data as a noisy linear map *x*_*d*_ = *ax*_*b*_+*b*+*ξ*, where *a* = 1 was associated with the adder mechanism and *a* = 0 with the sizer, under the assumption of a fixed threshold. They reported *a ≃* 0.871, 1.02, and 1.08 for the three temperatures, placing the data closer to the adder regime. Susman *et al*. applied a fold-growth regression and found substantial lineageto-lineage heterogeneity, interpreted as variation in the implemented control mechanisms, with an average close to adder behavior [19]. Finally, the data in [16] were also reported to be broadly consistent with adder-like behavior. [16].

To reconcile this we note that an adder-like regression slope does not uniquely identify an adder mechanism when the division threshold fluctuates with temporal correlations [12]. As a concrete demonstration, we show in Appendix Section A-A that under a sizer division control mechanism, with threshold correlation *ρ*_*u*_, the noisy linear-map slope is approximately *a ≃* 2*ρ*_*u*_. A sizer mechanism can therefore produce a slope *a ≃* 1 when *ρ*_*u*_ *≃* 1*/*2, giving the appearance of an adder in the above interpretation. In contrast, the likelihood framework developed here does not assume a-priori the typical timescale of the threshold, and therefore no such ambiguity arises.

Despite having a statistically significant winner in most datasets, we find that the likelihood gap among candidate mechanisms is generally small in all cases; the bars in Fig. 3 (top) are of similar height. The small likelihood gap can be seen also in the per-cell cumulative likelihood distributions, estimated for each candidate mechanisms over the pooled data from all experiments, presented in Fig. 3 (bottom). It implies the data occupy a regime in which all candidate mechanisms are quantitatively competitive and only weakly distinguishable. This is precisely the regime anticipated in the likelihood-gap discussion of Section II, and mirrors the simulation results: even with the ground-truth mechanism known to be one of the simple model mechanisms, others may still assign relatively high likelihood to the observations. This motivates us next to go beyond the previously studied models and to inquire whether the data contains structure not described by any of them.

### C. A data-driven division control rule learned from experimental data

In light of the small likelihood gap between previously suggested models, as revealed by our method, a natural question is whether a data-driven approach can identify a division control mechanism that better describes the data than any of those considered so far. To address this, we trained a flexible neural-network model directly on the lineage data. It receives current and previous-cycle cell-state variables as input, and learns the conditional distribution of cycle durations without being restricted to any specific control rule (see Materials and Methods Section IV-D for details). The average likelihood of each data-set given the rule learned by the neural network, is depicted as red bars in Figure 3 (top), showing that it consistently outperforms the other models. Its distribution of likelihood across cycles, from all data-sets pooled together (with balanced amounts from each experiment), is plotted as a red line in Figure 3 (bottom). Here, also, it achieves the highest average and highest likelihood throughout the majority of the range. These results suggest that the experimental data contain reproducible structure that is not fully captured by any one of the simplified model division rules. What is this structure, and can we find an approximation to the data-driven division control rule?

Having learned the conditional distribution of cycle durations from the data, we characterize its dependence on cell state through the conditional mode, which represents the *most likely* cycle duration,

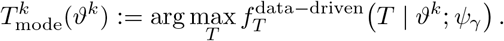

For the generalized-Gamma model used here, the mode has a closed-form expression (see Materials and Methods Section IV-D), allowing us to evaluate the most likely duration predicted by the model for each cell state.

Having obtained the precise modal prediction 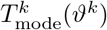 for each cell, we then seek a simple analytical approximation to the learned rule. Among a range of low-dimensional candidates, we found that the combination

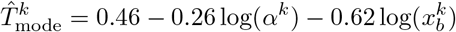

provides an excellent approximation to the network output, with Pearson correlation^1^ 0.864 with the exact modal prediction of the network. Thus, despite the high dimensionality of the input space, the learned control rule is captured to a remarkable degree by a simple affine combination of log(*α*) and log(*x*_*b*_). In comparison, the sizer combination 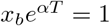, has a correlation value of 0.019 with the network output and the adder combination, 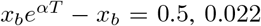 These comparisons show that the learned rule is substantially closer to a logarithmic affine law in (*α, x*_*b*_) than to either the sizer or adder laws.

Together, these results show that the learned model not only outperforms the simplified candidate mechanisms, but also reveals how the most likely cycle duration depends on the cell state. Although one could regress the observed durations directly on the cell-state variables, such an analysis would

^1^The Pearson correlation between two series *x*^*k*^ and *y*^*k*^ is,

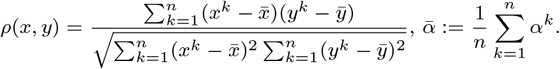

be intrinsically noisy because each observation is only a single realization from its conditional distribution. In addition, standard regression typically estimates the conditional mean, which differs substantially from the mode for the asymmetric, long-right-tailed distributions relevant here. Learning the full conditional distribution instead provides direct access to the most likely duration and reveals a low-dimensional timing law that cannot be recovered as reliably from the raw observations alone or summarized by the sizer, adder, or LFC relations. A full visualization of the model is provided in Appendix Section G; interpretation of the approximation above is outside the scope of this work and and is left for future research with more extensive data analysis.

## III. Discussion

This work reformulates division-mechanism identification as a maximum-likelihood problem and thereby replaces heuristic classification with a rigorously defined quantitative criterion. Rather than asking whether lineage data appear visually or phenomenologically closer to one candidate mechanism or another, we ask which model assigns higher likelihood to the data. This constitutes a change of perspective, providing both a quantitative measure for competing mechanisms, and a general tool for future application to new classes of candidate control mechanisms.

A key theoretical result is that, within the model class considered here, discrimination among division mechanisms is determined by the conditional distribution of cell-cycle durations. Other aspects of the lineage model, including growth-rate variability and division asymmetry, affect the full data-generating process but not the likelihood-ratio comparison among mechanisms.

An important aspect of the theory is its modularity. The likelihood formulation separates division-mechanism identification from the specification of the growth-rate model, so these two problems need not be solved jointly. In the present work, a particular stochastic growth model was sufficient to develop and evaluate the likelihood framework. However, future work can study richer and more detailed models of growth-rate dynamics separately, and then incorporate them into the broader lineage model without changing the logic of fair mechanism comparison, provided that their parameters remain independent of the mechanism-specific parameters. This separation opens the door to a more systematic research program in which improved growth models and improved division models can be developed in parallel.

Application of our approach to both simulations and experimental data paint a consistent picture: simple candidate models of cell-division control are statistically identifiable in principle but only weakly separated in practice. In simulations, the true generating mechanism is reliably assigned the highest likelihood, yet competing mechanisms remain quantitatively plausible, demonstrating that formal identifiability need not imply strong empirical separation. A similar pattern emerges across the six *Escherichia coli* datasets: the sizer, adder, and related models provide meaningful and competitive descriptions,but no single mechanism dominates across all datasets and growth conditions. Thus, the experimental data do not support a sharp, universal choice among the simplified control rules considered here. The consistently higher likelihood achieved by the flexible data-driven model indicates that the lineage data contain additional structure not fully represented by any single simplified models.

More broadly, the framework introduced here provides a foundation for cumulative progress in the study of stochastic division control. New mechanistic hypotheses can now be evaluated on a common, quantitative likelihood scale. We expect this perspective to be useful not only for bacterial division control, but also for other biological systems in which hidden stochastic thresholds govern observable transitions.

## IV. Materials and Methods

Section II formulated mechanism identification in terms of the conditional likelihood of cell-cycle durations. The framework requires, for each mechanism *γ*, the ability to evaluate the conditional density 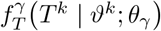 For the LFC mechanism, this density admits an analytical expression, and its likelihood can therefore be evaluated directly, see Appendix Section A-C.

The situation is different for the sizer and adder mechanisms. In both cases, the cycle duration is defined implicitly through a threshold-crossing event, and after the specialization introduced in Section II, the required likelihood term is a first-passage-time density of the form

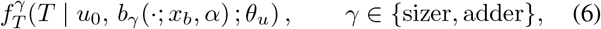

a special case of (2). For the Ornstein–Uhlenbeck threshold model (A.2) with a time-dependent barrier, no closed-form expression for (6) is available in the regimes relevant here [5], [8], [10], [11], [15], [28]. As a result, the sizer and adder likelihoods cannot be evaluated analytically. To obtain a closed-form and computationally efficient approximation to these likelihood terms, we proceed in two stages. First, we empirically determine a parametric family that accurately approximates simulated first-passage-time distributions (FPTD). Second, we train neural networks to map cycle-specific inputs to the parameters of that family.

### A. Empirical identification of a parametric family for simulated FPTDs

For a configuration (*u*_0_, *x*_*b*_, *α, θ*_*u*_), we generate *M* i.i.d. sample 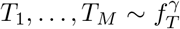 that form the target first-passage-time distribution (*u*_0_ is the previous cell-size for sizer or the previous cell added-size for adder). Figure 4 illustrates this procedure for a single configuration; See Proposition B.1 in the Appendix for a closed-form solution and transition law of the OU threshold, enabling efficient simulations.

**Fig. 4.**
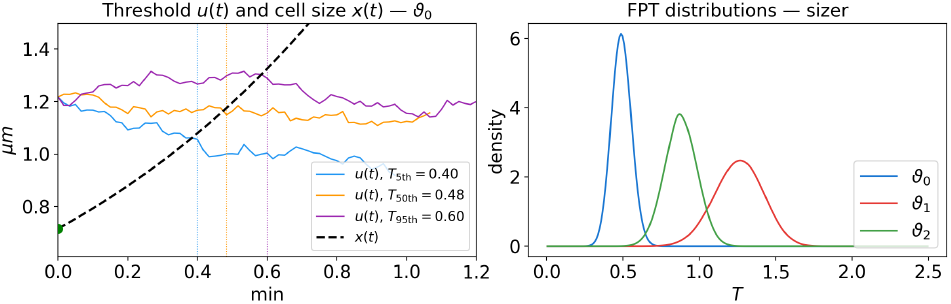
First-passage-time distributions. Three parameter configurations *ϑ*_0_, *ϑ*_1_, *ϑ*_2_ are selected with growth rate *α* = 1 and OU mean *μ*^*u*^ = 1 for all three; the remaining parameters are (*u*_0_, *x*_*b*_, *σ*_*u*_, *τ*_*u*_) = (1.22, 0.72, 0.12, 2.1), (0.96, 0.29, 0.26, 6.7), and (0.92, 0.39, 0.16, 3.8). **(Right)** Empirical FPT distributions for the three configurations, estimated from 10^5^ simulated samples each. The three distributions are visually distinct, illustrating the sensitivity of the FPT distribution to the parameters. **(Left)** For configuration *ϑ*_0_, three instantiations of the stochastic threshold process *u*(*t*) are shown, corresponding to FPTs at the 5th, 50th, and 95th percentiles of the simulated distribution. The deterministic cell-size trajectory 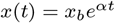 (dashed black) is shared across all realisations; division occurs at the first time *x*(*t*) crosses *u*(*t*).

We then ask which parametric family best approximates these empirical distributions. For each configuration (i.e. M samples), we fit several candidate distributions and find that the generalized-gamma family consistently provided the best approximation, see Appendix Section B. We therefore adopt it as the surrogate family for the sizer and adder first-passage-time distributions. We use the three-parameter formula

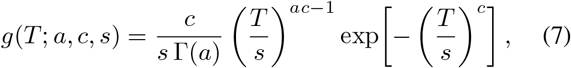

with shape parameters *a >* 0, *c >* 0, and scale parameter *s >* 0. This family is sufficiently flexible to capture the skewness, tail behavior, and modality of the simulated first-passage-time distributions observed under both mechanisms.

### B Neural-nets surrogates for sizer and adder

Having identified the generalized-gamma family as a suitable parametric approximation for the simulated first-passage-time distributions, we train a separate neural surrogate for each of the sizer and adder mechanisms. Fix a mechanism *γ ∈ {*sizer, adder*}* and a parameter configuration

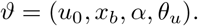

The goal is to learn the map

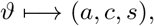

where (*a, c, s*) are the parameters of the generalized-gamma density in (7). The surrogate is trained by minimizing the negative log-likelihood of the simulated first-passage times under the predicted generalized-gamma density. Once trained, the surrogate provides an explicit closed-form approximation to the otherwise unavailable first-passage-time likelihood term,

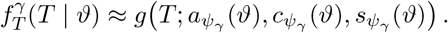

Substituting it into the likelihood formulation of Section II yields tractable, differentiable, and computationally efficient likelihoods for both the sizer and adder mechanisms. We also trained baseline linear regressions which achieved significantly degraded results, justifying the use of a neural network. Appendix Section D provides full details on the regression and the neural network training.

### C. Dimensionless representation and practical likelihood evaluation

To enable a unified analysis across datasets, we work with a unit-free representation obtained by deterministic normalization within each dataset. This normalization serves two purposes. First, it removes dataset-specific size and time scales, allowing the likelihood-based analysis to be carried out in a common coordinate system across datasets. Second, it reduces the dynamic range of the variables entering the surrogate models, thereby improving numerical stability.

For a given dataset, let 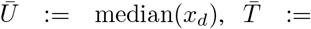 median(*T*), denote empirical size and time normalizations respectively, where the medians are computed over all cells in the dataset. We then define the normalized variables

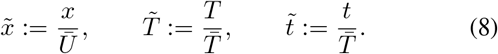

Accordingly, the per-cycle growth rate is transformed to the dimensionless quantity 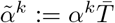

The transformation 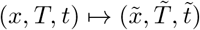 is deterministic and invertible. Consequently, any likelihood comparison carried out in the normalized coordinates is equivalent to the corresponding comparison in physical units up to a fixed change of variables. In particular, identifiability conclusions are invariant under this normalization. Throughout the manuscript each dataset has been transformed to this normalized representation and therefore we omitted the tildes from the notation.

The likelihood-ratio formulation in 4 extends naturally from a single lineage to a complete dataset. In real world data, a small number of cell cycles may be corrupted by segmentation errors, tracking mismatches, or rare biological events that are not adequately captured by the model. Because the dataset likelihood is a product of per-cell likelihood terms, even a few such outlying observations can drive the overall likelihood to an artificially small value and thereby dominate the optimization. To reduce this sensitivity, we use a robust, model-adaptive trimming procedure known as *trimmed maximum-likelihood* [4], [6], [13], see Appendix Section C.

### D. A data-driven generalized-gamma model learned from real data

We train a separate neural model directly on real lineage data in order to learn a data-driven division rule unconstrained by any canonical mechanism. Three datasets come from the MC4100 strain measured in the mother machine at three temperatures, 37^*°*^C, 27^*°*^C, and 25^*°*^C, as reported by Tanouchi *et al*. [21], [22]. These experiments provide long single-cell trajectories recorded at one-minute resolution and include, respectively, 160, 54, and 65 mother-cell lineages after processing [21]. Two additional datasets come from the study of Sassi *et al*. [16], who measured *E. coli* cells in a mother-machine device under two nutrient conditions, denoted as *rich* and *poor*. The sixth dataset, denoted here as *Salman MG1655*, comes from the long-trace mother-machine measurements of wild-type MG1655 cells reported by Susman *et al*. [19].

To this end, we concatenate all lineages from the six real datasets after converting each dataset separately to the dimensionless representation (8). Because the datasets differ in size, a naive concatenation would overweight the largest datasets. We therefore balance the training set so that each dataset contributes a comparable number of cells to the optimization procedure.

Here, the configuration 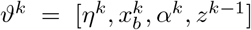 includes both current-cycle and all previous-cycle information. The representation provides a flexible, feature-rich description of the cell state while remaining agnostic to any specific canonical division mechanism. As in the synthetic surrogate setting, the real-data network, parameterized by a weight vector *Ψ*_*γ*_, outputs the parameters (*a, c, s*) of a generalized-gamma density and is trained by minimizing the negative log-likelihood of the observed normalized cycle durations where we denote *γ* = data driven for the data-driven mechanism. For the asymmetric first-passage-time distributions relevant here the most likely value is the conditional mode, given for a generalized-gamma density by,

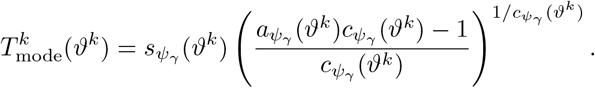

This model is not constrained to represent a sizer, an adder, or an LFC mechanism. Instead, it directly learns from the real data which covariates best predict the full conditional distribution of cycle durations. The role of this network is therefore complementary to that of the canonical-mechanism likelihoods. The sizer, adder, and LFC models test specific mechanistic hypotheses, whereas the real-data generalized-gamma network provides a flexible empirical benchmark that reveals how well an unconstrained distributional model can explain the data. See Appendix Section E for complete details on the neural network architecture and training procedure.

The procedure developed in this section provides the conditional cycle-duration likelihoods required by Section II. Sections II-A, II-B and II-C uses these learned and analytical likelihoods for two complementary purposes. First, to test identifiability on simulated data by comparing the optimized likelihoods of the canonical mechanisms; and second, to analyze six real datasets, where the canonical models are compared against the flexible data-driven model and the learned data-driven network is subsequently interpreted analytically and visually. Code that reproduces our results and allows, using our trained models, inference of other datasets is available at https://github.com/RonTeichner/newBacteria.

## Acknowledgment

The authors thank Kuheli Biswas and Omri Barak for valuable discussions.

## APPENDIX A

### Likelihood-based hypothesis testing

Our *inferential goal* as defined in Section II is to determine the division indicator *γ* from the observations *z*^0:*N−*1^, while treating the parameters *θ*_*γ*_ and *θ*_*x*_ as unknown. To that end, in Equation (3), we defined the likelihood under each hypothesis *γ* as

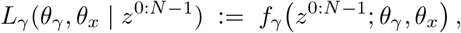

where for integers *I ≤ j*, we write *z*^*i*:*j*^ := *{z*^*i*^, *z*^*i*+1^, …, *z*^*j*^ *}*. Here we present a generalization of the results of Section II. We denote the *history* available at cycle *k* as *H*^*k*^ := *z*^0:*k−*1^. Let

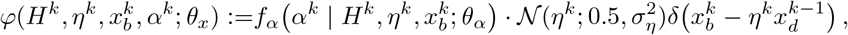

be a factor that collects all terms associated with growth-rate variability, division asymmetry, and the deterministic mass-conservation recursion. By construction, it depends on *θ*_*x*_ but not on *θ*_*γ*_.

#### Proposition A.1

*Under the model family defined in Section II, the likelihood under mechanism γ factorizes as*

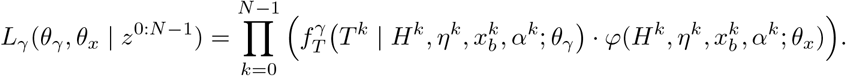

Proposition A.1 expresses the likelihood as a product of two conceptually distinct factors - a mechanism-dependent term involving the conditional density of cycle durations, and a mechanism-independent term involving growth-rate variability and division asymmetry.

To compare two candidate mechanisms *γ* and *δ*, we defined the generalized likelihood ratio in Eq. (4). Under standard regularity conditions for non-nested, possibly misspecified models, the generalized likelihood-ratio statistic is asymptotically optimal in that it consistently selects the mechanism closest to the true data-generating process in Kullback–Leibler divergence [24]; because no finite-sample optimality is guaranteed in this general setting, we therefore assess robustness by a subsampling-based bootstrap procedure, repeatedly re-estimating the maximum-likelihood solutions on randomly thinned datasets and reporting the resulting mean and variability across runs.

The next Lemma is a generalization of Eq. (5) in the main text.

#### Lemma A.2

*Consider any two division mechanisms γ and δ. Then the generalized likelihood ratio* (4) *reduces to*

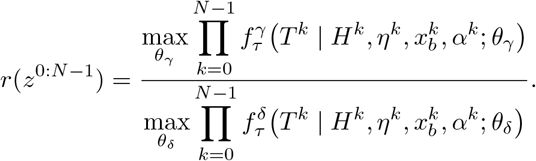

*In particular, the nuisance parameters θ*_*x*_ *cancel from the likelihood ratio*.

Lemma A.2 shows that, within this broad family, discrimination among division mechanisms is determined solely by the likelihood of the observed cycle durations. Growth-rate variability and division asymmetry affect the full data-generating model, but do not affect the generalized likelihood ratio as long as their parameters are independent of *θ*_*γ*_. The proofs of Proposition A.1 and Lemma A.2 are at Appendix Section A-B.

#### A. Misclassification as adder by the noisy linear map with a correlated threshold

While in the current work we classify between division-control mechanisms via Lemma A.2, in a previous analyses classification was performed using the noisy linear map

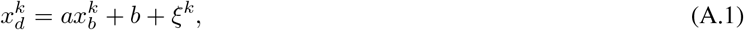

where *a ≃* 0 is conventionally associated with a sizer and *a* 1 with an adder [22]. Here we show why this interpretation need not hold when the division threshold is temporally correlated.

Consider first the sizer mechanism. Division in cycle *k* occurs when

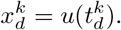

Because the birth size is inherited from the preceding division,

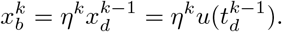

For clarity, suppose initially that division is symmetric, *η*^*k*^ = 1*/*2, and denote

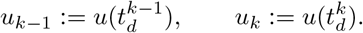

It then follows that

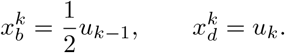

To expose the effect of threshold memory, approximate the OU latent threshold process (see Appendix A-C) transition between consecutive division events by

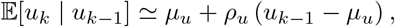

as in Proposition B.1, where *ρ*_*u*_ denotes the effective correlation of the threshold between consecutive division events. Substituting 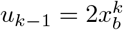 gives

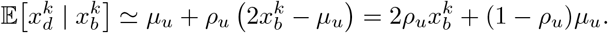

Comparison with Eq. (A.1) yields

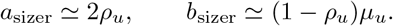

The heuristically expected sizer result *a ≃* 0 is therefore recovered only when threshold memory between successive divisions is negligible, *ρ*_*u*_ *≃* 0. In contrast, for a value of 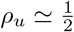, the sizer mechanism would produce

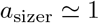

and would consequently be classified as an adder by the linear-map criterion.

#### B. Proofs

*Proof of Proposition A*.*1*. Recall that the likelihood of an observed lineage under each hypothesis *γ* is

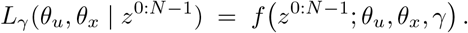

Since *η*^0^ is not measured we assume without loss of generality *η*^0^ = 0.5 and therefore, for mathematical convenience we set 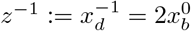 The causal structure implies that the likelihood factorizes as

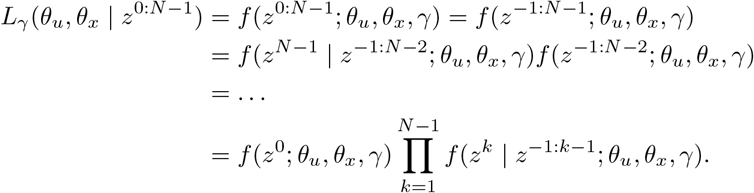

The second term further factorizes as,

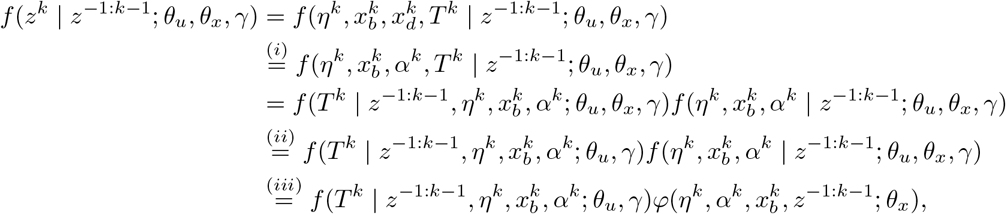

where we defined,

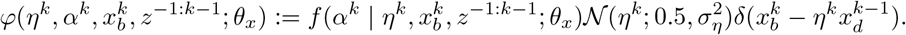

In the derivation above, (*i*) is because the model assumes perfect exponential growth by which 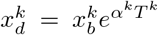, and so 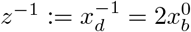 in (*ii*) we dropped *θ*_*x*_ because we conditioned on all the variables it affects, and (*iii*) results from

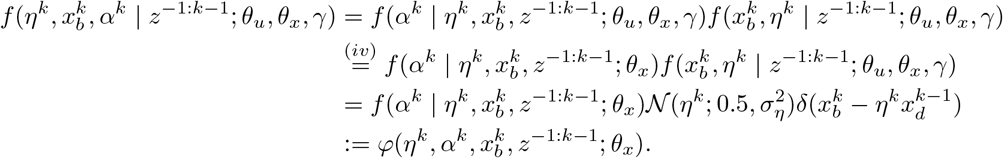

Here (*iv*) results from our model assumption by which the PDF of *α* is independent on the threshold parameters.Regarding the term corresponding to the first cell in the lineage, *f* (*z*^*−*1:*N−*1^; *θ*_*u*_, *θ*_*x*_, *γ*), it factorizes as

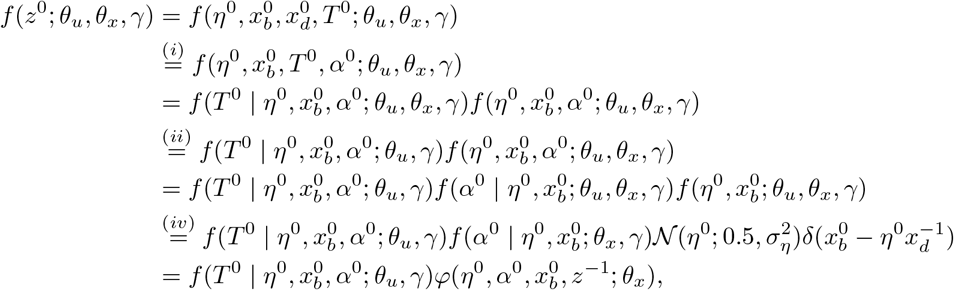

where the justifications for (*i*), (*ii*), (*iv*) are as above. Thus, the complete likelihood is given by,

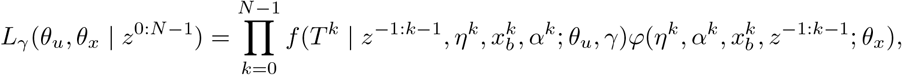

and this completes the proof..

*Proof of Lemma A*.*2*. Consider any two division mechanisms *γ* and *δ*. Substituting the likelihood from Proposition A.1 into the likelihood ratio (4) and noticing that the likelihood is a product of functions either of *θ*_*x*_ or of *θ*_*u*_ we obtain,

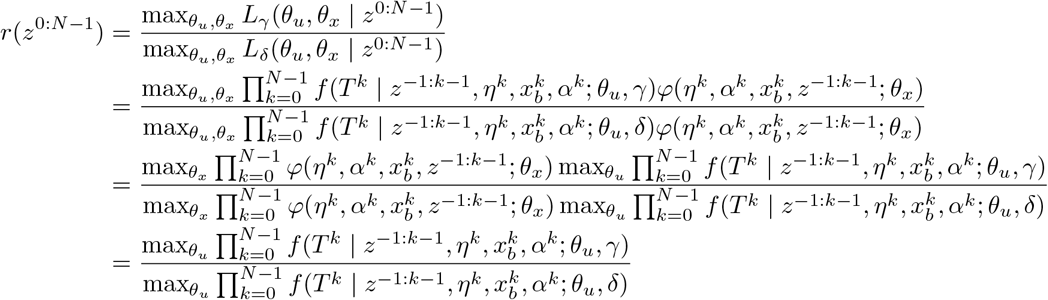

and this completes the proof.

#### C. Special case mechanisms

Candidate hypotheses that have been suggested include a *sizer*, where division is triggered upon reaching a critical size, and an *adder*, where division occurs after a fixed size increment has been added over the cycle [1], [9]. We also consider a *log-fold-change* (LFC) mechanism, in which division is governed by a target multiplicative increase over the cycle [2]. We now show that these models are all contained in the broad family above.

##### a)Sizer and adder as first-passage-time models

For the sizer and adder hypotheses, we introduce a latent threshold process *u*(*t*) modeled as an Ornstein–Uhlenbeck (OU) process,

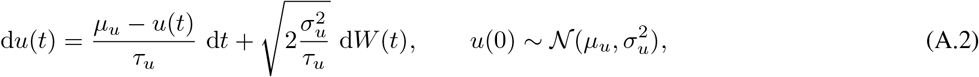

with parameters *θ*_*u*_ = [*μ*_*u*_, *τ*_*u*_, *σ*_*u*_]. Under the *sizer* mechanism, division occurs when the cell size reaches the threshold, *x*(*t*) = *u*(*t*). Under the *adder* mechanism, division occurs when the added size since birth reaches the threshold, 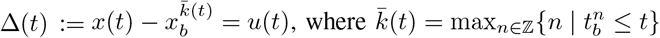 denotes the current cycle index.

##### FPT survival and density

Let *s*(*t*) be a continuous Markov process with *s*(0) = *s*_0_, and let *b*(*t*) be a (deterministic) barrier with *b*(0) *< s*_0_. The first-passage time to the barrier is

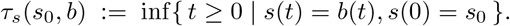

Its survival function is *F*_*τ*_*s* (*t*; *s*_0_, *b*) = *P* (*τ*_*s*_(*s*_0_, *b*) *> t*) and the corresponding FPT density (FPTD) is

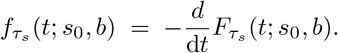

In our setting, *s*(*t*) *≡ u*(*t*) is the OU process (A.2), *s*_0_ is the threshold value at cycle start, and *b*(*t*) *≡ b*_*γ*_(*t*; *x*_*b*_, *α*),

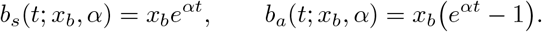

Hence, for *γ ∈ {*sizer, adder*}*, (2) takes the form

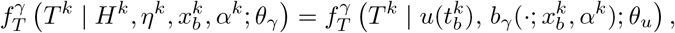

the probability-density of observing a cycle length *T*^*k*^ when the threshold starts at 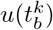, the barrier parameters are 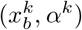 and the OU parameters are *θ*_*u*_.

##### c)Log-fold-change (LFC)

Under the LFC mechanism, the division threshold is taken to be constant within each cycle but random across cycles,

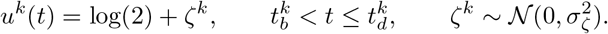

Division occurs when this threshold is first met by the barrier

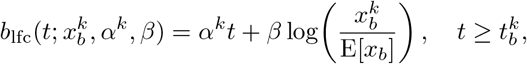

where *β* is a mechanism parameter and E[*x*_*b*_] denotes the mean birth size in the relevant dataset [2]. Since the barrier is linear in time and the threshold is cyclewise constant, the first-passage time is obtained explicitly as 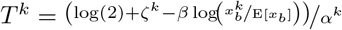 Therefore, with *θ*_lfc_ := [*β, σ*_*ζ*_], the conditional density of the cycle duration is

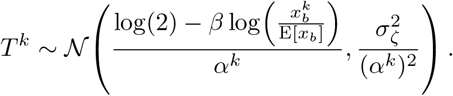

Thus, sizer, adder, and LFC all belong to the general family through appropriate specifications of (2); for LFC the resulting conditional cycle-duration density is available analytically, whereas for sizer and adder Appendix B introduces a methodology for learning corresponding closed-form surrogate densities.

## APPENDIX B

### Selection of a parametric surrogate family for simulated first-passage-time distributions

To obtain a tractable surrogate for the simulated first-passage-time distributions (FPTDs) of the sizer and adder mechanisms, we first carried out a systematic parametric distribution-selection procedure. The goal was to identify a low-dimensional family that provides an accurate approximation to the empirical FPTDs generated by the stochastic-threshold simulations over a broad range of parameter settings.

#### 1) Simulated FPT samples

Fix a mechanism *γ ∈ {*sizer, adder*}* and a parameter configuration

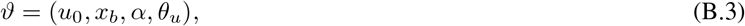

where *θ*_*u*_ denotes the mechanism-specific threshold parameters and (*u*_0_, *x*_*b*_, *α*) collects the cycle-specific variables required to define the corresponding first-passage problem. See Section B-5 for details on sampling configurations. For each such configuration, the simulator generates i.i.d. samples of first-passage times (see Proposition B.1 below)

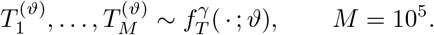

These samples define the empirical cumulative distribution function (CDF)

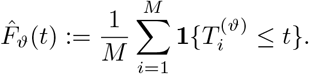

##### Proposition B.1

(Closed-form solution and transition law of the OU threshold). *Let u*(*t*) *satisfy the Ornstein–Uhlenbeck SDE* (A.2) *with parameters θ*_*u*_ = [*μ*_*u*_, *τ*_*u*_, *σ*_*u*_] *and initial condition u*(0) = *u*_0_. *Then, for every t ≥* 0,

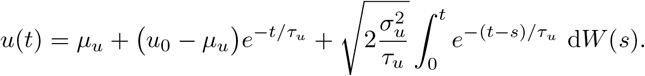

*(See, e*.*g*., *[15]*.*) In particular, conditional on u*(0) = *u*_0_, *the random variable u*(*t*) *is Gaussian with*

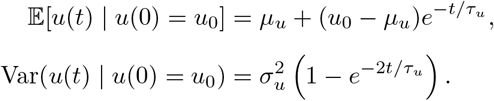

#### 2) Parametric fitting of candidate families

Let 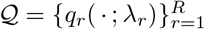denote the collection of candidate parametric families considered in the analysis, where *q*_*r*_(; *λ*_*r*_) is a density on (0, *∞*) indexed by parameter vector *λ*_*r*_ *∈* Λ_*r*_. For each mechanism *γ ∈ {*sizer, adder *}*, each sampled parameter configuration *ϑ*, and each candidate family *q*_*r*_, we estimate the fitted parameter vector by maximum likelihood,

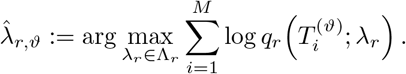

This yields the fitted density

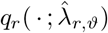

and its associated fitted CDF

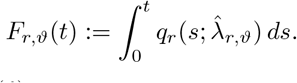

Thus, for every simulated FPT sample-set 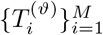, each candidate parametric family is fitted independently and produces one fitted distribution *F*_*r,ϑ*_.

#### 3) Wasserstein–1 comparison with the empirical FPTD

To evaluate the quality of fit, we compare each fitted distribution *F*_*r,ϑ*_ to the empirical FPT sample-set 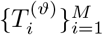 using the one-dimensional Wasserstein–1 distance,

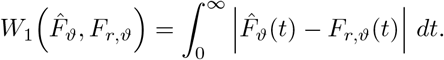

This formula makes clear that *W*_1_ measures the integrated discrepancy between the empirical and fitted cumulative distributions. In the implementation, this comparison is evaluated numerically from the simulated sample and the fitted distribution. The Wasserstein–1 distance is chosen because it captures distributional differences in both shape and support, making it more robust to misalignments than KL divergence [14]. This formulation captures the total “mass” that must be moved to transform one histogram into the other, weighted by the distance it is moved.

**Fig. B.1.**
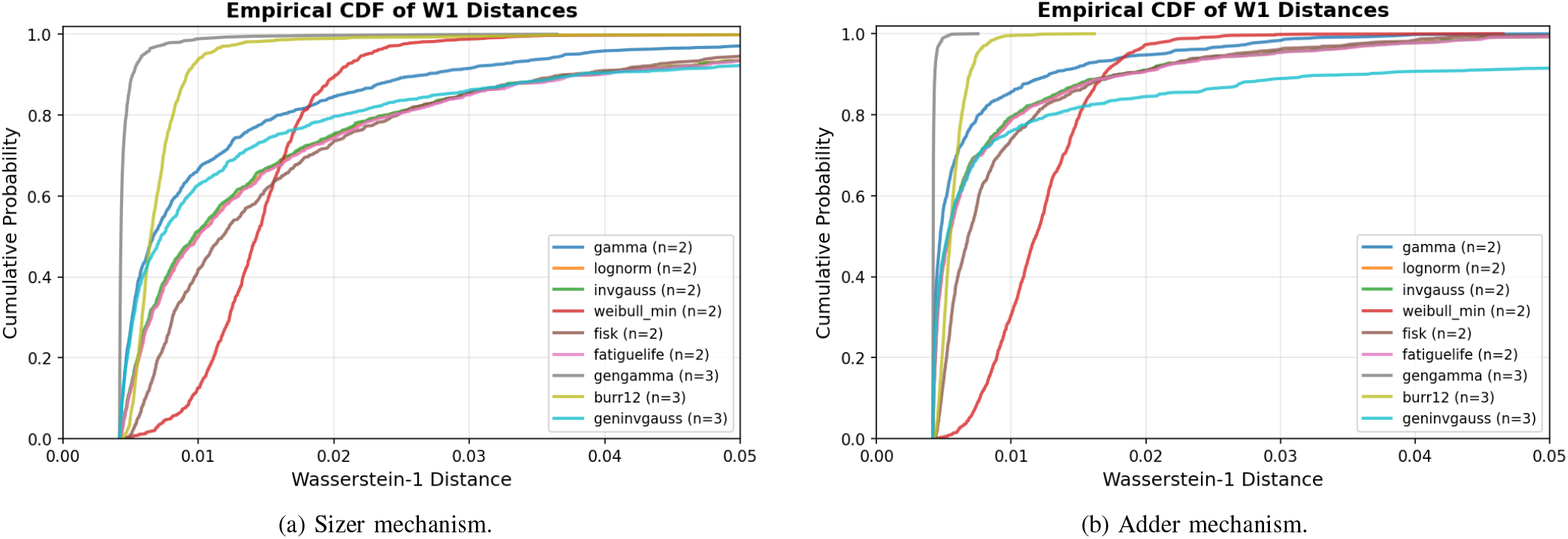
Wasserstein–1 distance between the empirical FPT distributions obtained from simulation and the fitted candidate parametric distributions, namely Gamma, log-normal, inverse Gaussian, Weibull, log-logistic (Fisk), Birnbaum–Saunders, generalized gamma, Burr XII, generalized inverse Gaussian, and a two-component Gaussian mixture. Panel (a) corresponds to the sizer mechanism and panel (b) to the adder mechanism. Lower values indicate a more accurate approximation to the empirical first-passage-time distribution. Across the sampled parameter settings, the generalized-gamma family provides the best overall fit and was therefore selected as the parametric surrogate family used in the main text. Since we work in dimensionless units, the units of the Wasserstein–1 distance are cell cycles.

For each candidate family *q*_*r*_, the distribution-selection analysis is repeated over many sampled parameter settings *ϑ*_1_, *…, ϑ*_*S*_. The overall performance of family *q*_*r*_ is then summarized by aggregating the corresponding Wasserstein distances, through the empirical mean

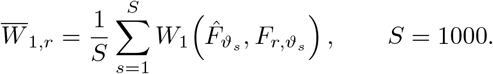

The candidate families are then ranked according to these aggregated Wasserstein discrepancies.

#### 4) Outcome of the selection analysis

The comparison revealed that the generalized-gamma family (see Eq. (7) in the main text) consistently provided the best overall approximation to the empirical FPTDs of both the sizer and adder mechanisms. In particular, it achieved the smallest Wasserstein–1 discrepancies across the sampled parameter range, indicating that it captures the skewness, tail behavior, and modality of the simulated FPT distributions more accurately than the alternative candidate families. Figure B.1 summarizes the Wasserstein–1 comparison between the fitted candidate distributions and the empirical first-passage-time histograms.

This empirical model-selection step motivated the surrogate-learning strategy adopted in the main text-instead of approximating the FPTD directly by a high-dimensional histogram, we train a neural network to output the parameters of a generalized-gamma density, and then optimize that network by negative log-likelihood on the simulated first-passage times.

1. *Sampling the configuration space for synthetic FPTDs:* Rather than drawing complete configurations, *ϑ* in (B.3), from a fixed joint model, we sample each variable independently of the others. This intentionally enlarges the support of the training set and exposes the surrogate model to a broader range of combinations than would arise under a more restrictive generative prior. The goal is not to mimic any one biological dataset exactly, but rather to ensure that the learned surrogate *f*^*γ*^ remains

accurate throughout the range of parameter values encountered in practice.

The growth rate *α* is sampled from the empirical histogram of *α*-values observed in the real datasets, after transforming the data to dimensionless variables. This preserves the experimentally observed range and shape of the growth-rate distribution while avoiding the need to impose a specific parametric form at this stage.

For the sizer mechanism, we sample the threshold parameters according to

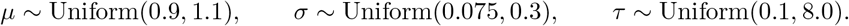

Here, *μ* is not fixed exactly at 1, even in dimensionless units, because the normalization is performed using empirical medians rather than exact means; allowing *μ* to vary in a narrow interval around 1 accounts for a possible shift between these two reference scales. The ranges for *σ* and *τ* are deliberately broad and were chosen to include, and somewhat exceed, the support of the corresponding values observed in real data.

Conditional on (*μ, τ, σ*), we then sample the initial threshold value according to

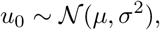

and the birth size according to

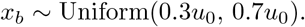

Here, the range is chosen to include the support of division asymmetry observed in the real datasets.

For the adder mechanism, we begin from the same sampled configuration and then apply the transformation

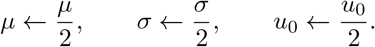

Notably, the correlation time *τ* is left unchanged, and the sampled birth size *x*_*b*_ is also left unchanged. This transformation reflects the fact that, in the adder formulation, the relevant threshold and initial offset are measured in added-size units rather than absolute-size units, while the temporal scale of threshold fluctuations remains the same.

## APPENDIX C

### Practical optimization via model-adaptive trimming

Let 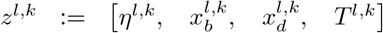 denote the k-th observed cell cycle in lineage l, and let 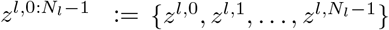denote the full set of observations from lineage *l*, where *N*_*l*_ is the number of observed cycles in that lineage. The complete dataset is then 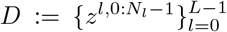, where *L* is the number of lineages. Assuming that the lineages are conditionally independent given *θ*_*γ*_, the generalized likelihood ratio between two candidate mechanisms *γ* and *δ* becomes

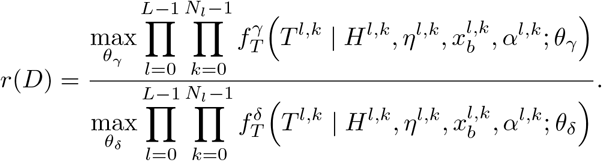

That is, the dataset-level comparison is obtained by multiplying the per-cell conditional cycle-duration likelihoods across all observed cycles in all lineages.

In real world data, a small number of cell cycles may be corrupted by segmentation errors, tracking mismatches, or rare biological events that are not adequately captured by the model. Because the dataset likelihood is a product of per-cell likelihood terms, even a few such outlying observations can drive the overall likelihood to an artificially small value and thereby dominate the optimization. To reduce this sensitivity, we use a robust, model-adaptive trimming procedure known as *trimmed maximum-likelihood* [4], [6], *[13]. For a fixed mechanism γ* and parameter value *θ*_*γ*_, consider the collection of per-cell likelihoods

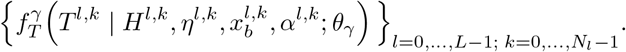

Let *q*_*ρ,γ*_(*θ*_*γ*_; *D*) denote the empirical *ρ*-quantile of this collection, where 0*≤ρ <* 1 (we will often set *ρ* = 0.15). We then retain only those cells whose likelihood exceeds this threshold, namely

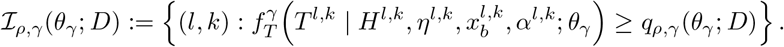

The resulting trimmed likelihood is defined by

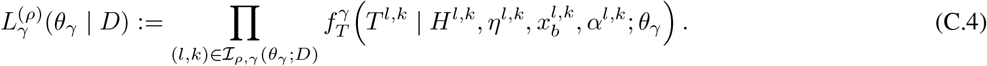

Thus, for each candidate parameter value, the model itself determines which observations are treated as low-likelihood outliers, rather than requiring these cells to be specified in advance. Accordingly, the dataset-level generalized likelihood ratio is replaced by its trimmed version, 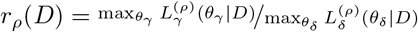 In other words, mechanism comparison is performed using only the highest-likelihood fraction 1 *− ρ* of cells under each candidate mechanism and parameter value.

## APPENDIX D

### Training of the generalized-gamma surrogates for sizer and adder

This appendix describes the training procedure used to construct the generalized-gamma surrogate likelihoods for the sizer and adder mechanisms. The neural surrogate is trained by combining a supervised warm start on fitted generalized-gamma parameters with a direct likelihood-based fine-tuning stage on raw first-passage-time (FPT) samples. The same script also trains a sparse affine baseline using Lasso regression with a one-standard-error (1SE) selection rule. Each configuration is represented by the six basic variables

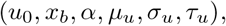

and the learning algorithms augment these by their logarithms, yielding the 12-dimensional input vector

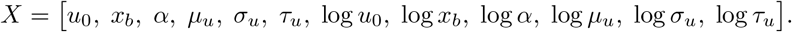

The targets for the neural surrogate are the generalized-gamma parameters in log-space (see Eq. (7) in the main text),

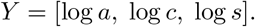

The logarithmic parameterization guarantees positivity of *a, c*, and *s* after exponentiation. The configurations are split into training and validation sets using a random train/validation split with 20% validation. Both the inputs *X* and the targets *Y* are standardized with the scalers fit on the training set only and then applied to the validation set.

The surrogate is a three-head multilayer perceptron. The network consists of a shared trunk followed by three scalar output heads corresponding to log *a*, log *c*, and log *s*.

*a)Shared trunk*.: Given input dimension 12, the trunk applies a sequence of fully connected blocks of the form

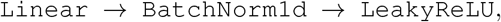

with optional dropout after each hidden block. The hidden-layer widths are (128, 128, 64), and the dropout probability is 0.02.

*b)Output heads*.: Each of the three heads receives the shared representation and applies the sequence

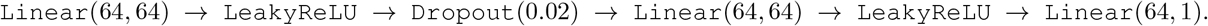

Thus, each head has two hidden layer of width 64 before the final scalar output, and the full network predicts the three log-parameters jointly.

*C)Phase 1: supervised warm start by mean-squared error:* Training begins with a supervised warm start on the fitted generalized-gamma parameters. Let 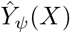 denote the network output in standardized target coordinates. The warm-start objective is the mean-squared error

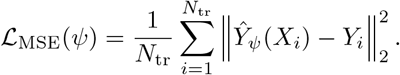

The model is optimized using the Adam optimizer with lr = 10^*−*3^, weight_decay = 10^*−*5^. Mini-batches are drawn with batch size of 64. After each gradient step, gradients are clipped to norm 1.0. Validation MSE is evaluated after every epoch, and the model checkpoint with the smallest validation loss is retained. This warm-start phase runs for 50 epoch

d)*Phase 2: negative-log-likelihood fine-tuning on raw FPT samples:* After the warm start, the best phase-1 checkpoint is used to initialize a second phase in which the network is trained directly on raw FPT samples. For a mini-batch of configurations, the network predicts (log *a*, log *c*, log *s*), which are exponentiated to obtain (*a, c, s*). For each configuration in the batch 1000 FPTs are sampled from the simulator. These sampled times are concatenated across the batch, and the objective is to minimize the mean negative log-likelihood under the predicted generalized-gamma density

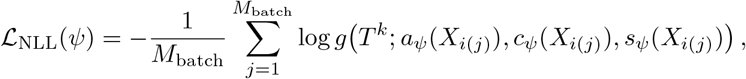

where *M*_batch_ is the total number of sampled FPTs in the batch and *i*(*j*) maps each sampled FPT to its parent configuration. Optimization again uses Adam, with a learning rate lr = 10^*−*3^ and the same weight decay 10^−5^. A cosine-annealing learning-rate scheduler is applied over the fine-tuning phase. Validation NLL is monitored after each epoch by evaluating the same generalized-gamma negative log-likelihood on subsampled validation FPTs. The best checkpoint is retained using early stopping. In addition to the neural surrogate, we train an affine model, to see if a neural model provides a substantial improvement.

**Fig. D.2.**
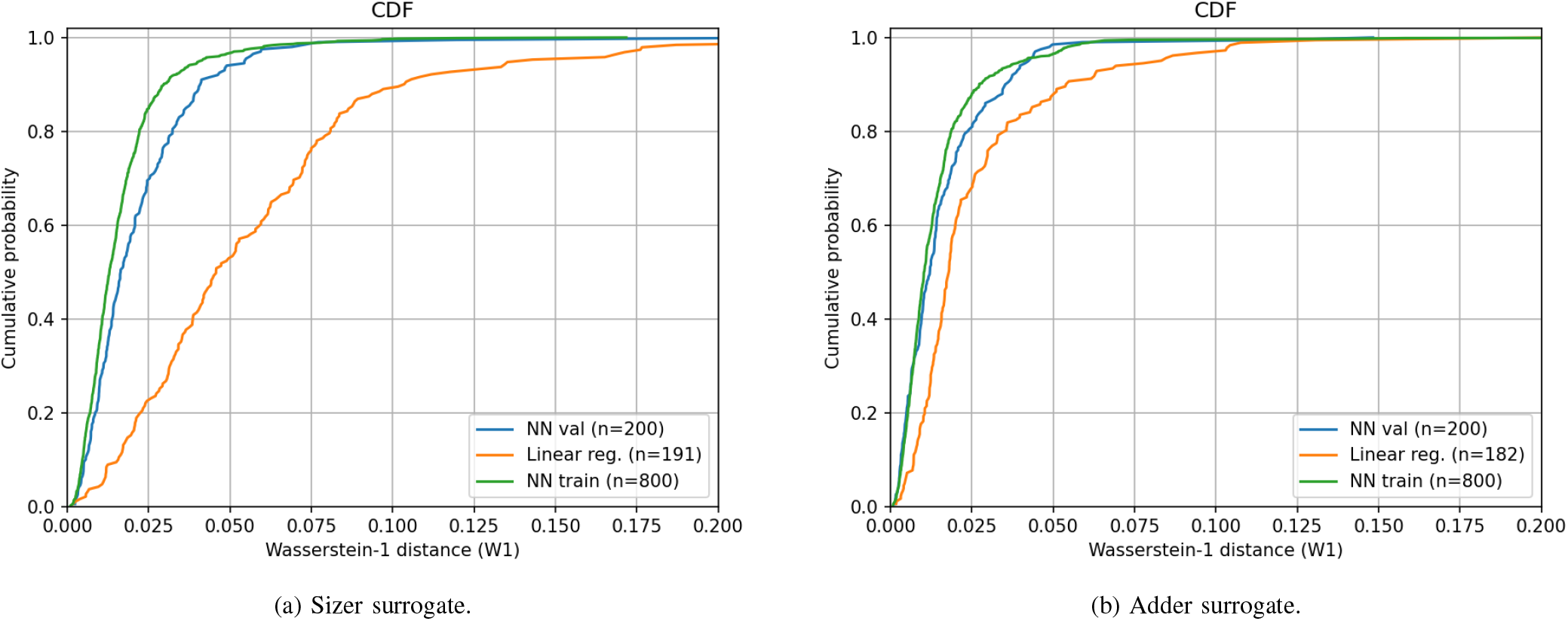
Wasserstein analysis of the generalized-gamma surrogate models. Panel (a) shows the validation analysis for the sizer surrogate, and panel (b) shows the corresponding analysis for the adder surrogate. In each case, the figure reports the empirical cumulative distribution of Wasserstein–1 distances between the true generalized-gamma fit and the generalized-gamma distribution predicted by the neural surrogate (green curve for training set and blue for validation set). The sparse affine Lasso model is also overlaid (orange curve), demonstrating the advantage of the neural surrogate.

1. *Validation in Wasserstein distance:* The validation figures for the surrogate models are generated as follows. For each configuration we draw samples from: 1)the *true* generalized-gamma distribution defined by the fitted parameters (*a, c, s*), see Section B-2, and 2)the *predicted* generalized-gamma distribution defined by the neural-network outputs (*a*_pred_, *c*_pred_, *s*_pred_).

The Wasserstein–1 distance between the two sampled distributions is then computed as the mean absolute difference between their sorted samples. The resulting collection of W1 distances across validation rows is summarized in an empirical cumulative distribution function (ECDF) of W1 values. In addition, the same figure overlays two additional curves:

1. the W1 distribution obtained from the sparse affine Lasso model, and
2. the W1 distribution of the neural surrogate on the *training* set.

Thus, the validation figure directly compares the neural surrogate, the sparse affine model, and the corresponding train-set performance in the same Wasserstein metric. Figure D.2a shows this analysis for the sizer surrogate, and Figure D.2b shows the corresponding analysis for the adder surrogate.

## APPENDIX E

### Generalized-gamma network trained directly on experimental data

This appendix describes the neural network used to learn a data-driven conditional distribution of cell-cycle durations directly from real lineage data. The training objective is a pure negative log-likelihood (NLL) on observed cell-cycle durations, after a warm-up stage based on linear regression.

a. *Input data and dataset normalization:* The train is performed on multiple datasets. Each dataset is normalized independently by dividing *x*_*b*_ and *x*_*d*_ by *U*_0_ = median(*x*_*d*_) and *T* by *T*_0_ = median(*T*), computed over all cells in that dataset.
b. *Per-cell feature construction:* After normalization, we flatten the lineages into per-cell training examples. Generation-0 cells are excluded, because the feature representation uses information from the previous cell cycle. For every cell, we construct a 14-dimensional feature vector

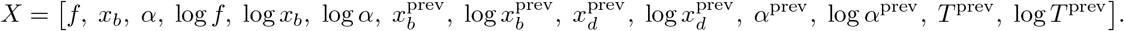

The prediction target for each row is the normalized cell-cycle duration *T*.
c. *Train/validation split and feature standardization:* All cells from all datasets are concatenated into a single pooled design matrix *X* and target vector *T*. We perform a random train/validation split using 20% for validation. Standardization is carried out by a fit on the training set only.
d. *Balanced sampling across datasets:* The datasets are balanced so that each dataset contributes equally during training and validation, regardless of its raw number of cells. Concretely, cells from dataset *i* are assigned weight proportional to the inverse of the number of training (or validation) cells from that dataset. This prevents the largest datasets from dominating the optimization purely through sample count.
e. *Network architecture:* The generalized-gamma real-data model uses the class MLP, which consists of a shared trunk followed by one scalar head per output dimension. For the generalized-gamma model, the output dimension is three, corresponding to

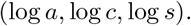

The shared trunk applies a sequence of blocks of the form

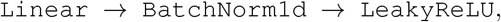

with optional dropout after each hidden layer. Each output head applies

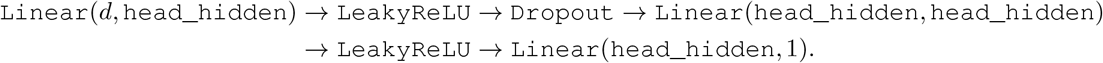

where *d* is the final shared hidden width. The script uses `head_hidden` = 32 for the real-data generalized-gamma model. The hidden widths are (128, 128, 64), with batch size 64, learning rate 10^*−*3^, dropout 0.0, patience 200, and trim fraction 0.85, see (C.4).

*f)Generalized-gamma likelihood:* The generalized-gamma log-density has an analytic term,

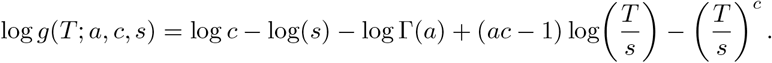

*g)Two-stage training procedure:* Training proceeds in two stages.

*h)Stage 1: MSE warm-up*.: Before likelihood optimization, we perform a warm-up stage in which the network is trained to match a linear-regression predictor of the observed cycle duration. Specifically, a `LinearRegression` model is fit to the standardized training inputs *X*_tr_ and the observed durations *T*_tr_. For each mini-batch, the network outputs are de-standardized and converted to approximate mean and scale surrogates. For the generalized-gamma model, we use

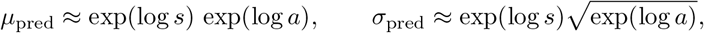

as simple warm-up proxies. These are compared against the linear-regression prediction 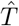 and a heuristic scale target 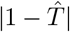 using a mean-squared error loss. This warm-up stage is intended only to anchor the network in a reasonable region of parameter space before the likelihood-based training begins.

*i)Stage 2: trimmed NLL training*.: After warm-up, the script performs the main optimization by minimizing the generalized-gamma negative log-likelihood on the observed durations. For each batch, the network predicts (log *a*, log *c*, log *s*), these are de-standardized and the per-cell negative log-likelihood values are computed. To reduce sensitivity to extreme outliers, the script applies trimming within each batch as defined in (C.4) - only the lowest-NLL fraction of cells is retained in the batch loss. The same trimming rule is also used during validation. This yields a robust likelihood criterion that prevents a small number of corrupted or highly atypical cells from dominating the objective.

*j)Optimizer, scheduler, and stopping rule:* Both training stages use the Adam optimizer. In the main likelihood-training stage, we additionally apply a cosine-annealing learning-rate schedule which monotonically reduces the learning rate across training. Validation NLL is monitored after every epoch, and the best network checkpoint is retained whenever the validation NLL improves by more than 10^*−*6^. Early stopping is triggered when the validation loss fails to improve for `patience` consecutive epochs. Thus, the final model is the checkpoint with the lowest validation NLL encountered during training, not necessarily the last epoch.

k)*Training-curve diagnostic:* Figure E.3 depicts the training diagnostic showing that the network is not in an overfitting regime.

**Fig. E.3.**
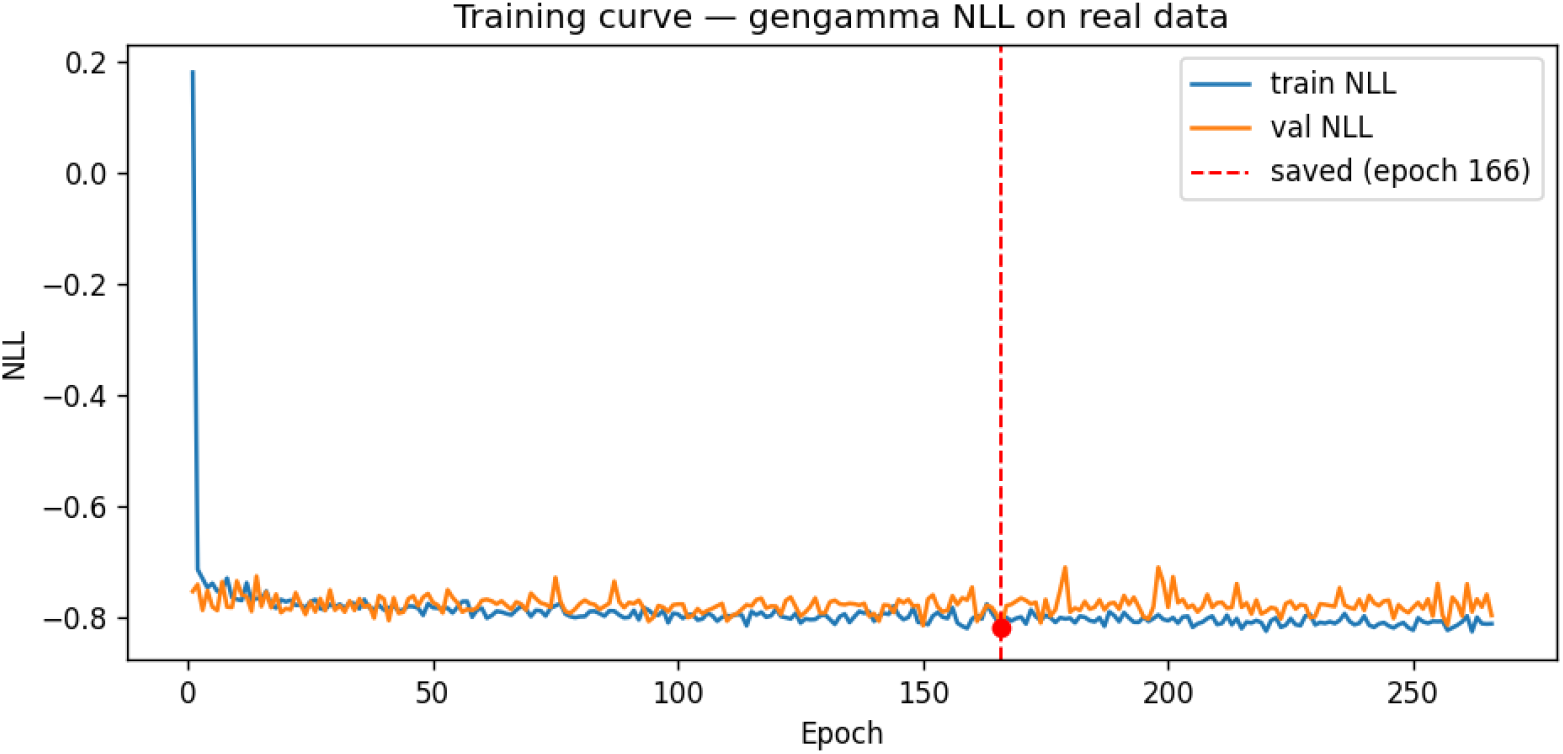
Training curve for the generalized-gamma network trained on real data. The figure plots the training and validation negative log-likelihood (NLL) as a function of epoch for the generalized-Gamma model fit directly to the pooled normalized experimental data. This diagnostic is used to assess convergence of the likelihood optimization and to select the best checkpoint under the early-stopping rule.

**Fig. F.4.**
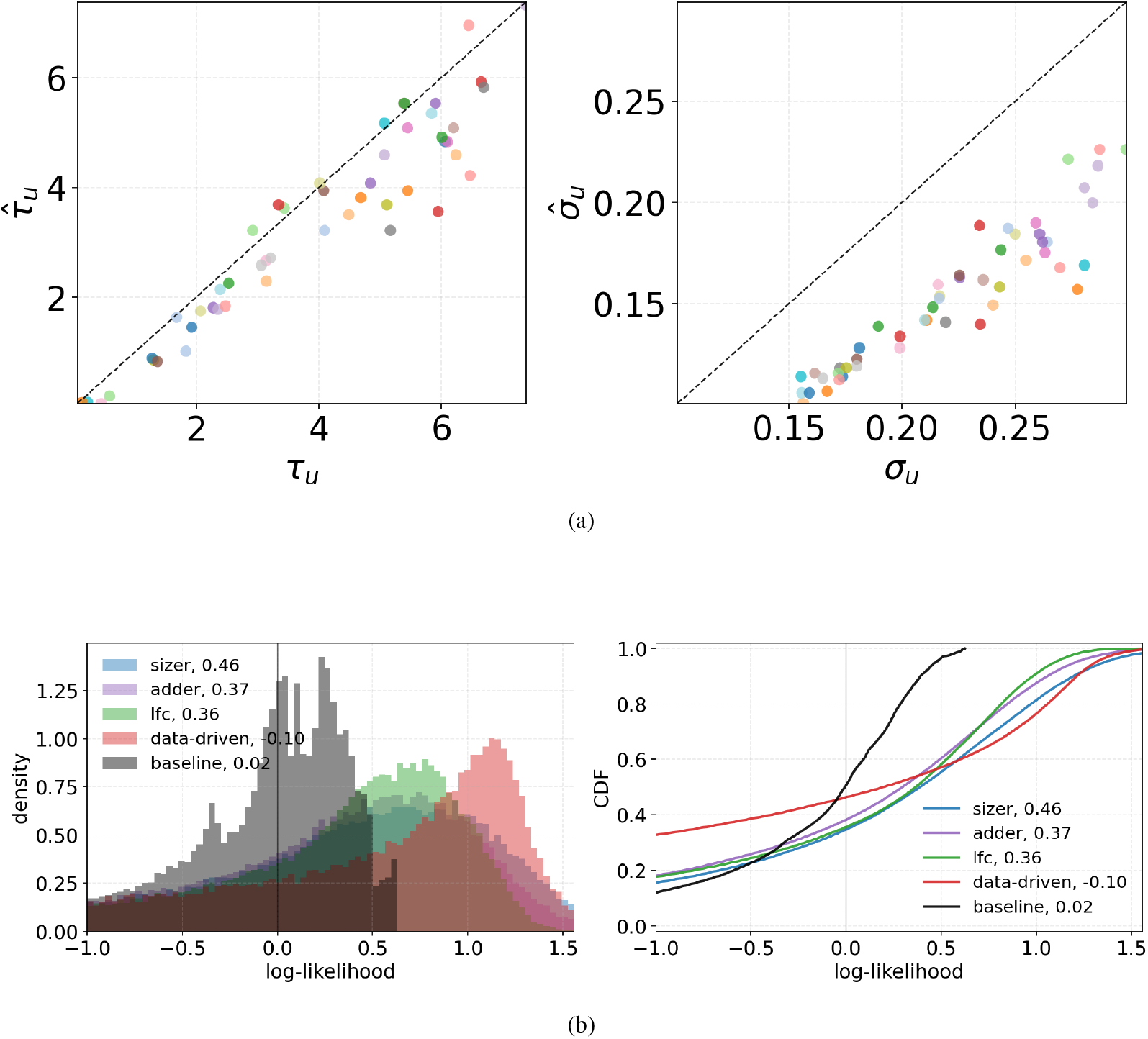
Additional simulation results for datasets generated under the sizer mechanism. Panel (a) shows dataset-level recovery of the generating parameters. The estimation of *μ*^*u*^ is degenerated due to normalization (see Materials and Methods Section B-5). A small systematic bias is visible, consistent with the classical finite-sample bias identified by Hurwicz [7] Panel (b) shows the pooled per-cell log-likelihood comparison across the candidate mechanisms and reference models.

## APPENDIX F

### Additional results

#### A. Additional simulation results

This appendix collects the full set of simulation figures for all three ground-truth mechanisms considered in this work: adder, sizer, and LFC. For each mechanism, we report two complementary diagnostics. The first compares the ground-truth parameter values used to generate the simulated datasets with the corresponding maximum-likelihood estimates, thereby assessing the accuracy of parameter recovery. The second summarizes the distribution of per-cell log-likelihoods under all competing models, namely sizer, adder, LFC, the data-driven FPTD model trained on real data, and a non-mechanistic baseline. Together, these figures provide a more complete picture of both parameter identifiability and model discrimination than the representative panels shown in the main text.

Figure 2 in the main-text summarizes the results for datasets generated under the adder mechanism and Figure F.4 presents the corresponding results for datasets generated under the sizer mechanism. Figure F.5 shows the simulation diagnostics for datasets generated under the LFC mechanism. In the main text we noted the fact that an incorrect mechanism may still provide a quantitatively good explanation of the data. This behavior is further illustrated here in Figure F.6, which shows that the sizer and adder mechanisms can achieve similar log-likelihood values at different parameter settings, underscoring that the incorrect mechanism may still explain the data well even though the true sizer mechanism is consistently favored. These results complete the simulation-based validation across the full set of canonical mechanisms studied in this work.

#### B. Additional experimental data results

Maximum-likelihood optimization was performed separately for each dataset and each mechanism. This produced one fitted parameter set per dataset and per mechanism, with the corresponding per-cell likelihood distributions and their statistics. The dataset-specific fitted parameter values are listed in Table F.1, and the statistics are given in Table F.2. The per-dataset normalization constants and lineage counts are reported in Table F.3. Figure 3 in the manuscript summarizes the grouped bar comparison of log-likelihoods and the pooled per-cell log-likelihood distributions across datasets, while Figure F.7 gives the detailed per-cell log-likelihood distributions for each individual dataset.

**Fig. F.5.**
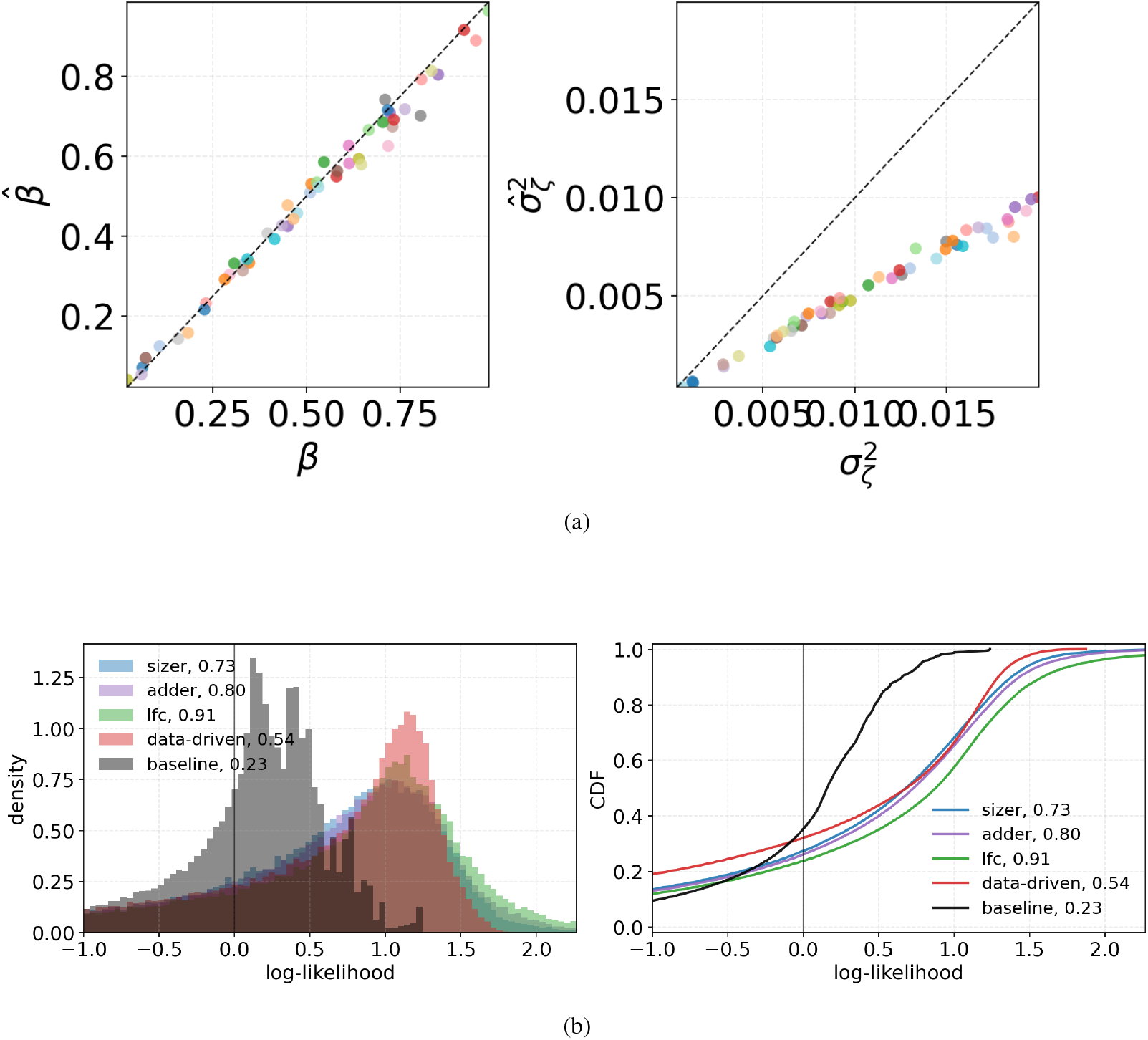
Additional simulation results for datasets generated under the LFC mechanism. Panel (a) shows dataset-level recovery of the generating parameters. A small systematic bias is visible, consistent with the classical finite-sample bias identified by Hurwicz [7]. Panel (b) shows the pooled per-cell log-likelihood comparison across the candidate mechanisms and reference models.

**Fig. F.6.**
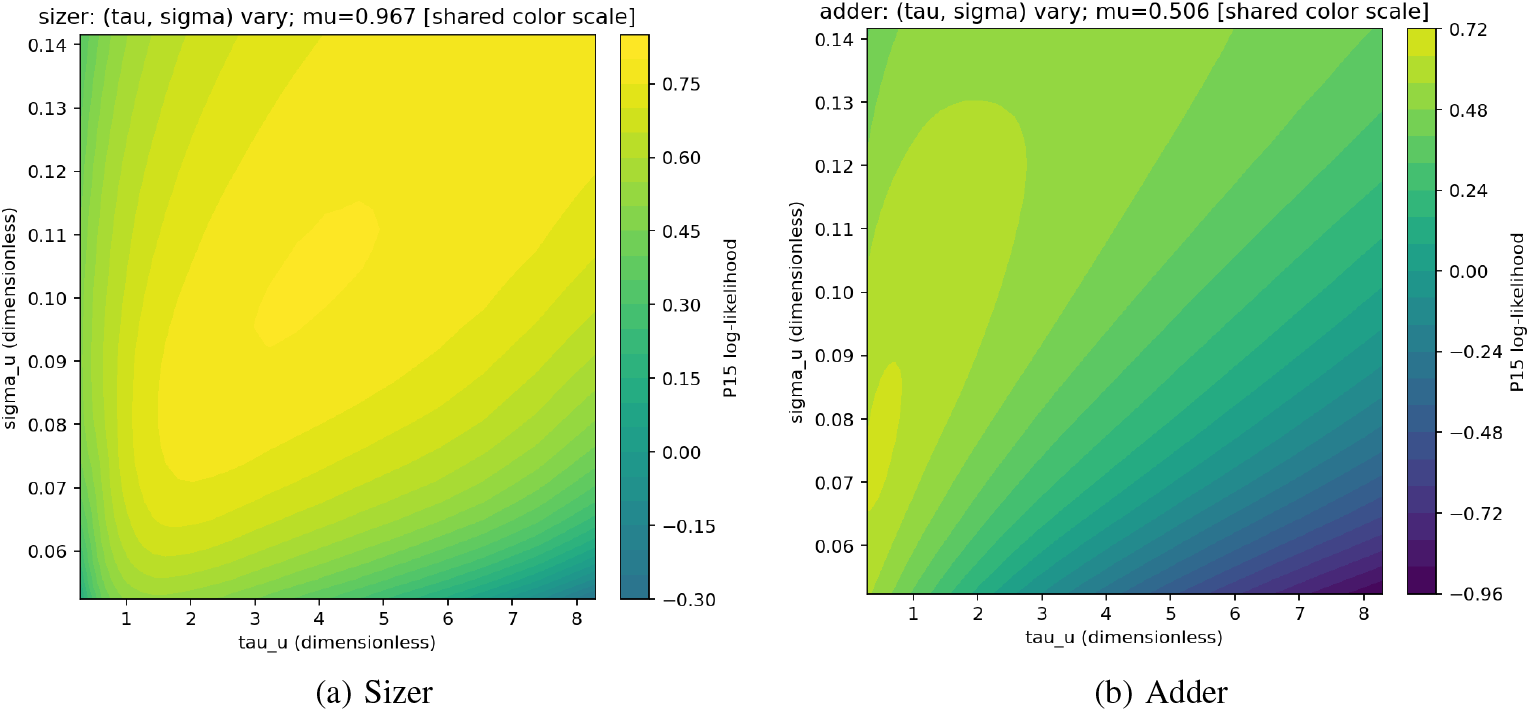
An adder model can achieve a likelihood similar to that of the generating sizer mechanism. Data were generated using the sizer mechanism, and their log-likelihood was evaluated under the sizer and adder hypotheses across their respective parameter spaces *θ*^*u*^. The heat maps show the resulting log-likelihood surfaces, computed using Eq. (C.4) with *ρ* = 0.15 (see Materials and Methods, Section IV-C). Although the sizer model achieves the maximum likelihood, a region of the adder parameter space yields nearly comparable likelihood values, demonstrating that an alternative division mechanism can remain quantitatively competitive.

**TABLE F.1.**
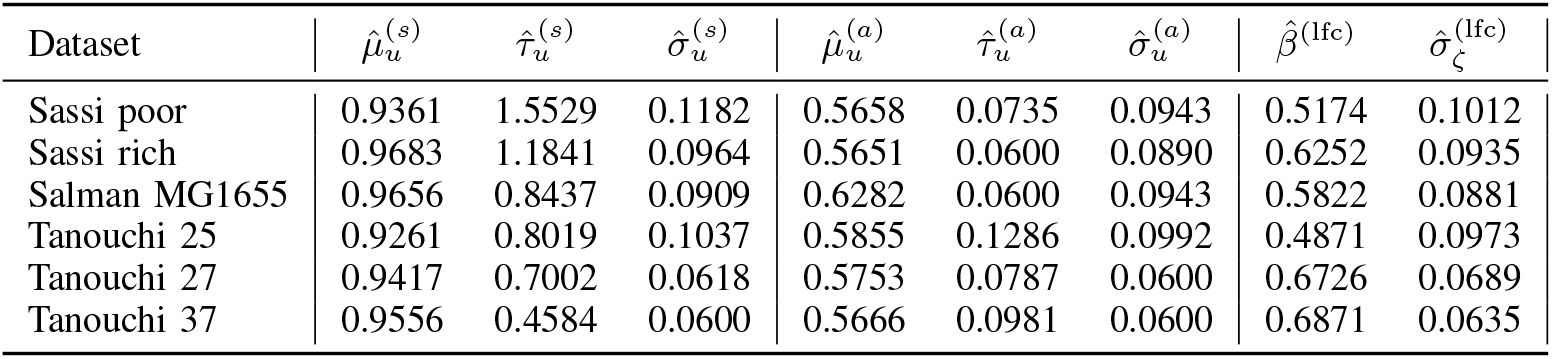
Dataset-specific maximum-lik elihood parameter estimates in the dataset-level analysis. In the dataset-level analysis, each dataset is fit separately under each mechanism (maximizing (c.4) with ρ = 0.15, see Materials and Methods section IV-C). The table reports the fitted threshold parameters for the sizer, adder, and LFC models.

**TABLE F.2.** Bootstrap mean log-likelihoods ± standard deviation in the dataset-level analysis.

| Dataset | $\ell_s$ | $\ell_a$ | $\ell_{\text{lfc}}$ | $\ell_{\text{data-driven}}$ | $\ell_{\text{base}}$ |
| --- | --- | --- | --- | --- | --- |
| Sassi poor | $0.4237 \pm 0.0214$ | $0.4378 \pm 0.0189$ | $0.4266 \pm 0.0222$ | $0.6070 \pm 0.0169$ | $-0.0641 \pm 0.0225$ |
| Sassi rich | $0.4982 \pm 0.0109$ | $0.4943 \pm 0.0087$ | $0.4486 \pm 0.0160$ | $0.6652 \pm 0.0082$ | $0.3924 \pm 0.0073$ |
| Salman MG1655 | $0.6133 \pm 0.0127$ | $0.5738 \pm 0.0119$ | $0.6316 \pm 0.0132$ | $0.6368 \pm 0.0142$ | $0.1644 \pm 0.0072$ |
| Tanouchi 25 | $0.6091 \pm 0.0112$ | $0.6019 \pm 0.0124$ | $0.6314 \pm 0.0109$ | $0.6664 \pm 0.0133$ | $0.1570 \pm 0.0100$ |
| Tanouchi 27 | $1.0677 \pm 0.0091$ | $1.0177 \pm 0.0070$ | $0.9920 \pm 0.0095$ | $1.1003 \pm 0.0114$ | $0.5835 \pm 0.0118$ |
| Tanouchi 37 | $1.0786 \pm 0.0043$ | $1.0410 \pm 0.0044$ | $1.0581 \pm 0.0041$ | $1.0902 \pm 0.0029$ | $0.5837 \pm 0.0038$ |

## APPENDIX G

### Full visualization of the learned data-driven fptd network

**TABLE F.3.** Dataset metadata for the dataset-level analysis. Per-dataset normalization constants and lineage counts extracted from `ml_per_dataset_results_real`Data_datasetLevel.csv.

| Dataset | $U_0$ | $T_0$ | $n_{\text{lineages, total}}$ | $n_{\text{lineages, used}}$ | $n_{\text{skipped}}$ |
| --- | --- | --- | --- | --- | --- |
| Sassi poor | 3.6835 | 80.0 | 219 | 218 | 1 |
| Sassi rich | 5.9394 | 45.0 | 92 | 92 | 0 |
| Salman MG1655 | 5.3374 | 18.0 | 20 | 20 | 0 |
| Tanouchi 25 | 4.1871 | 65.0 | 65 | 65 | 0 |
| Tanouchi 27 | 3.5947 | 53.0 | 54 | 54 | 0 |
| Tanouchi 37 | 4.4177 | 32.0 | 160 | 160 | 0 |

**Fig. F.7.**
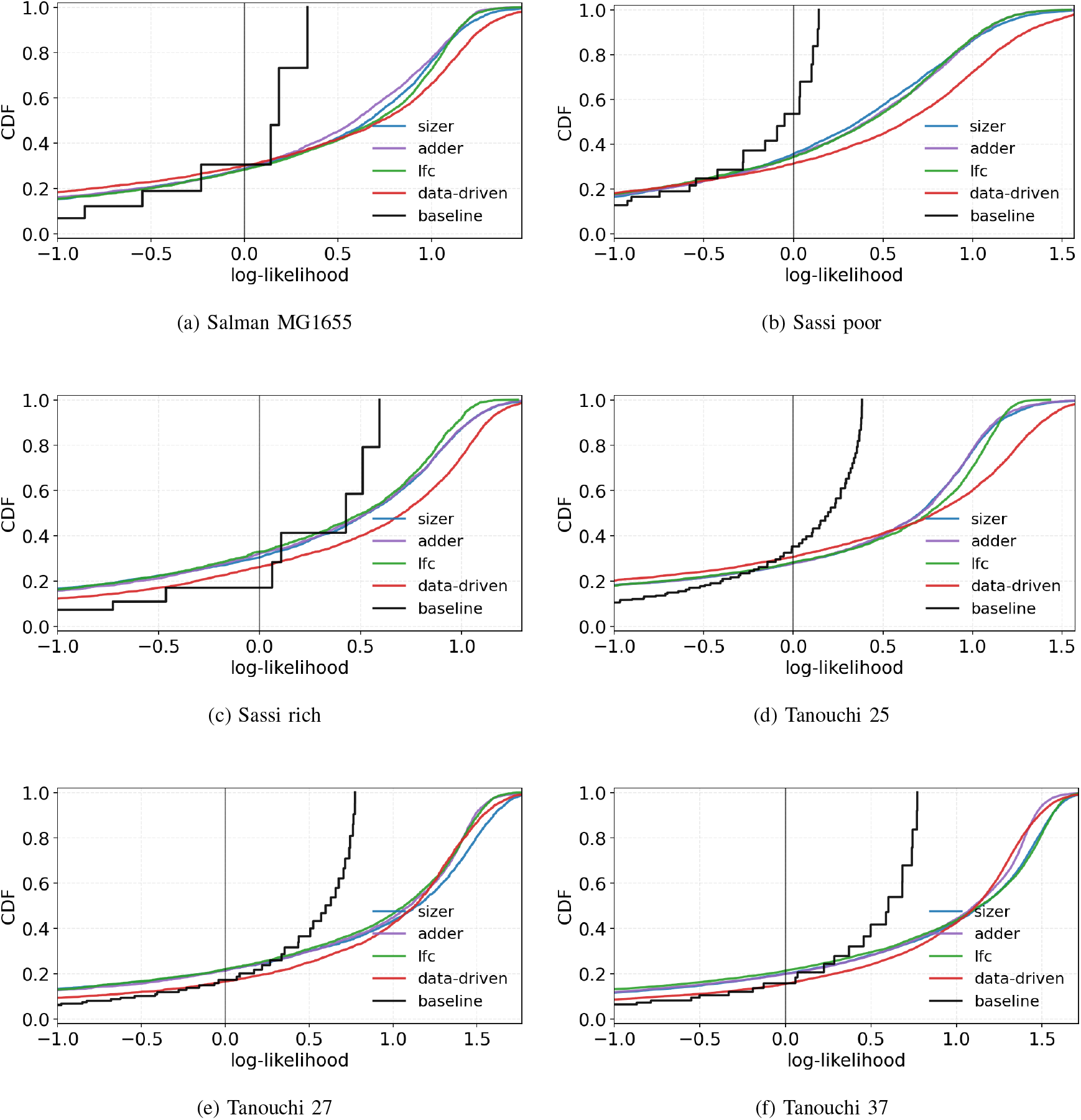
Per-dataset per-cell log-likelihood distributions for the dataset-level analysis. Each panel shows the per-cell log-likelihood comparison for one dataset, evaluated under the competing models after fitting each dataset separately. Together, these panels expose the dataset-specific shape of the per-cell likelihood distributions beyond the pooled summary shown in Fig. 3 in the manuscript.

**Fig. G.8.**
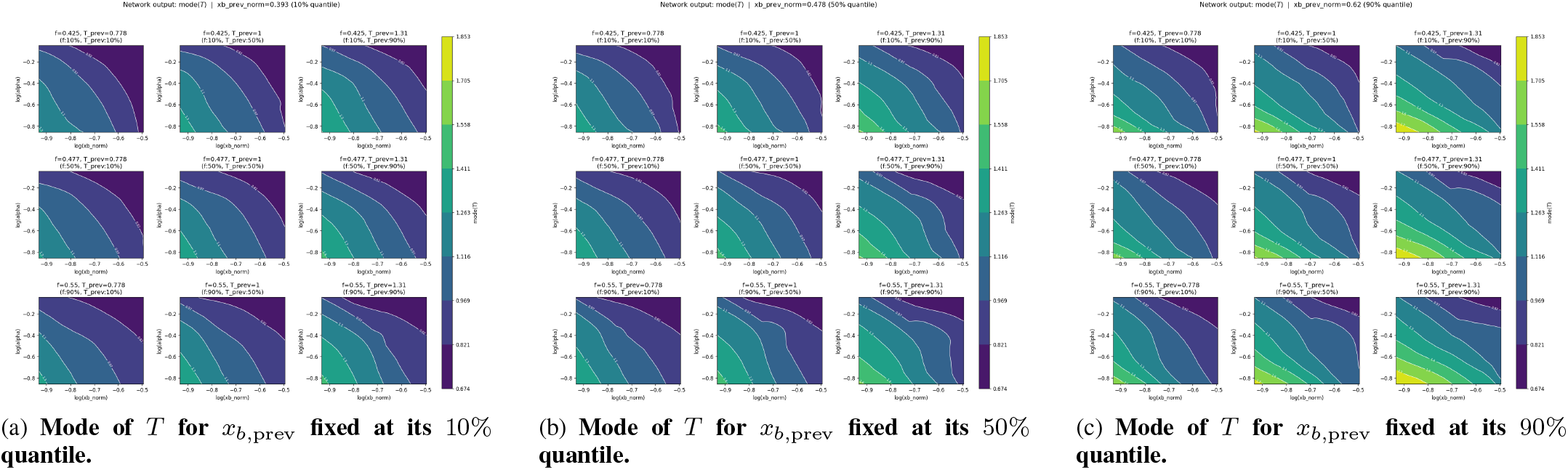
Mode of the learned cycle-duration distribution. Each panel is a 3 × 3 contour grid over (log *x*_*b*_, log *α*). Rows correspond to the 10%, 50%, and 90% quantiles of the current division fraction *f*, and columns correspond to the 10%, 50%, and 90% quantiles of the previous cycle duration *T*^prev^. Across the three panels, the previous birth size *x*_*b*,prev_ is fixed at its 10%, 50%, and 90% quantiles, respectively.

This appendix provides the full set of contour visualizations used to interpret the learned real-data generalized-gamma network. The figures are generated by evaluating the trained network on a structured grid over the two primary current-cycle variables, birth size *x*_*b*_ and growth rate *α*, while conditioning on representative values of additional covariates from the current and previous cycles. In all figures shown here, the horizontal axis is log(*x*_*b*_*/U*_0_), the vertical axis is log(*α*), the rows correspond to the 10%, 50%, and 90% quantiles of the current division fraction *f*, and the columns correspond to the 10%, 50%, and 90% quantiles of the previous cycle duration *T*_prev_. Each figure fixes the previous-cycle birth size *x*_*b*,prev_ at one of three representative quantiles: 10%, 50%, or 90%. Thus, every panel shows how the learned rule varies across the (*x*_*b*_, *α*) plane under a controlled choice of the remaining covariates.

The network outputs the generalized-gamma parameters (*a, c, s*) for each input state, from which the script derives interpretable statistics of the learned first-passage-time distribution. In the present appendix we focus on four such quantities: the mode of the cycle duration *T*, the coefficient of variation of *T*, the mean of *T*, and the variance of *T*. The mode is the most likely cycle duration under the learned conditional distribution, the coefficient of variation quantifies relative dispersion, and the mean and variance capture the central tendency and spread of the learned timing law. The mode figures correspond directly to the quantity used in Section II-C, while the remaining plots illustrate how the full learned distribution changes as a function of present and past cell-state variables.

1. *Mode of the learned cycle-duration distribution:* Figure G.8 shows the mode of the learned cycle-duration distribution,

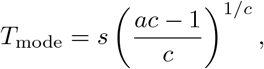

evaluated over the (log *x*_*b*_, log *α*) plane for three fixed quantiles of *x*_*b*,prev_. These plots provide the most direct visualization of the effective timing rule learned by the network. The comparison across panels shows how the modal prediction depends not only on the current-cycle variables *x*_*b*_ and *α*, but also on the previous-cycle state.
2. *Relative dispersion of the learned cycle-duration distribution:* Figure G.9 shows the coefficient of variation of the learned cycle-duration distribution,

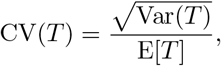

again across the (log *x*_*b*_, log *α*) plane and for the same three fixed quantiles of *x*_*b*,prev_. Because this quantity is strictly positive and can vary over a wide dynamic range, the script plots it using a logarithmic color scale. These figures show how the learned uncertainty in division timing changes across the observed state space and how it depends on previous-cycle covariates.
3. *Mean of the learned cycle-duration distribution:* Figure G.10 shows the conditional mean of the learned cycle-duration distribution,

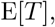

as a function of the current and previous cell-state variables. These plots complement the mode figures by showing how the center of the learned distribution changes across state space. Comparing Figures G.10 and G.8 illustrates the asymmetry of the learned distributions: the mean and the mode need not coincide, especially in regions where the learned distribution has a long tail.
4. *Variance of the learned cycle-duration distribution:* Finally, Figure G.11 shows the conditional variance of the learned cycle-duration distribution, Var(*T*), over the same set of fixed covariate quantiles. These plots quantify how the absolute uncertainty in division timing depends on the state of the present and previous cell cycles. In combination with Figures G.8–G.10, they provide a full visual picture of how the learned generalized-gamma rule changes both its location and its dispersion across the experimentally observed region of feature space.

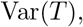

**Fig. G.9.**
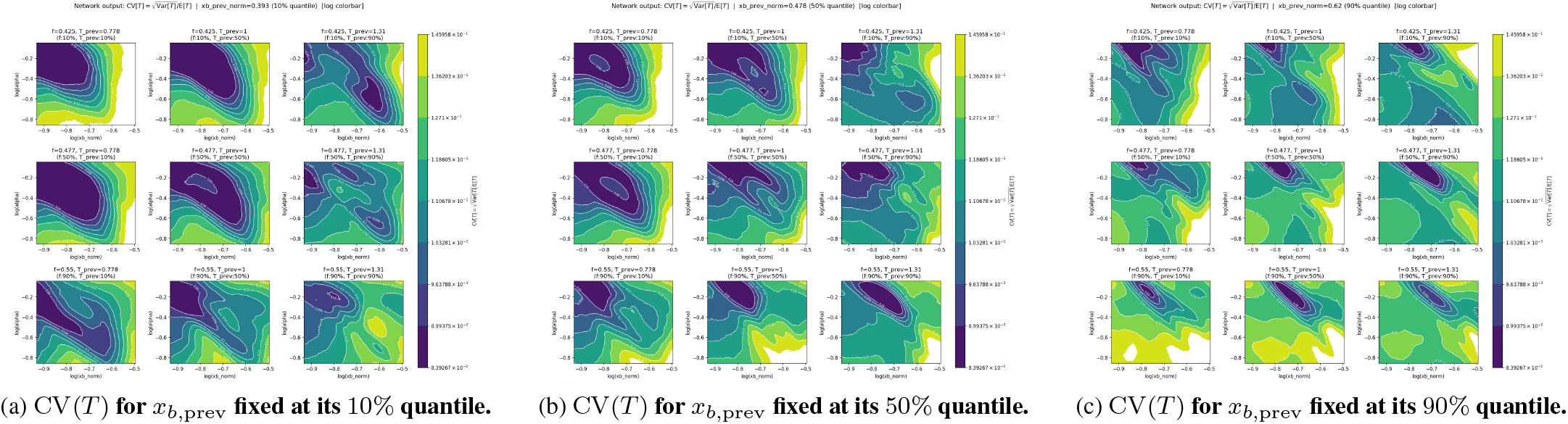
Coefficient of variation of the learned cycle-duration distribution. Each panel shows the network-derived CV(*T*) over the same 3×3 grid layout as in Fig. G.8. A logarithmic color scale is used in order to resolve the variation of CV(*T*) across the state space.

**Fig. G.10.**
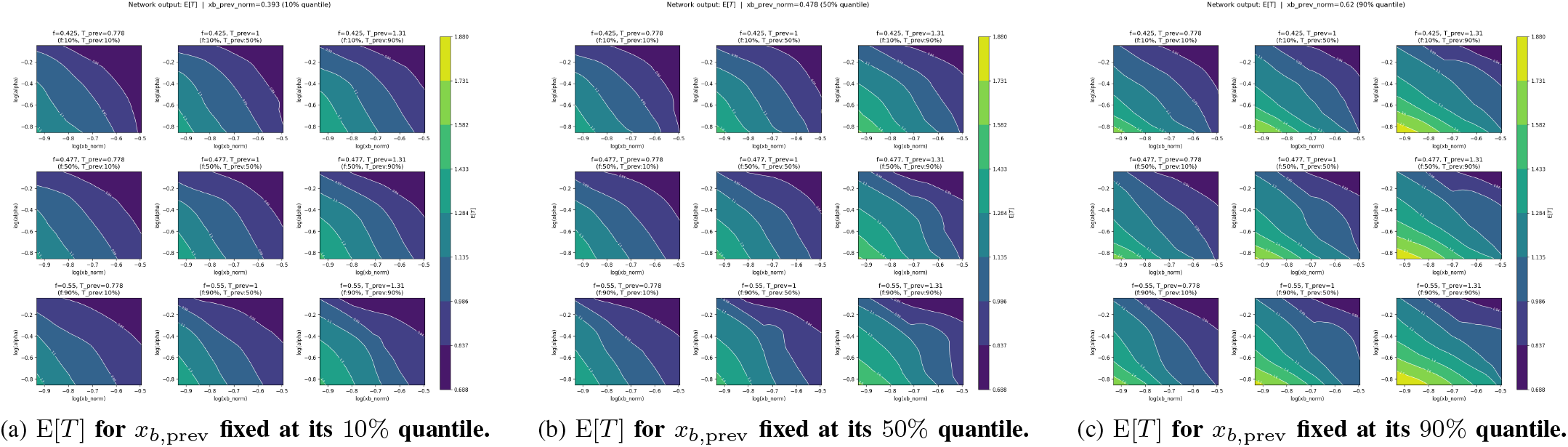
Mean of the learned cycle-duration distribution. Each panel shows the network-derived conditional mean E[*T*] over the same 3×3 quantile grid as in Fig. G.8.

**Fig. G.11.**
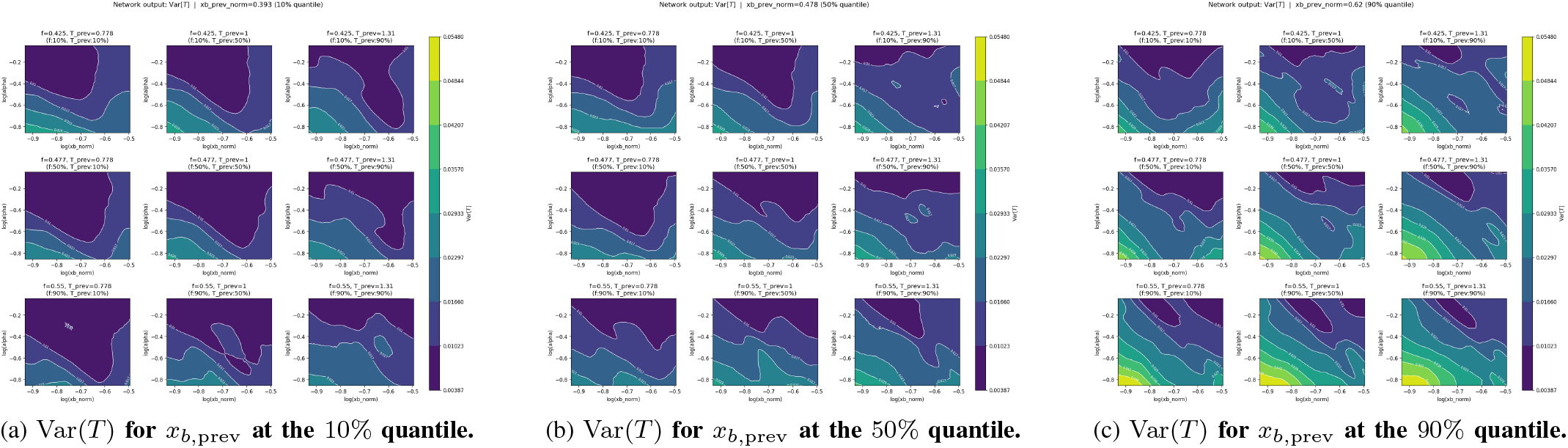
Variance of the learned cycle-duration distribution. Each panel shows the network-derived conditional variance Var(*T*) over the same 3× 3 quantile grid as in Fig. G.8.

## References

[1] Amir, A.: Cell size regulation in bacteria. Physical review letters 112(20), 208102 (2014)

[2] Biswas, K., Brenner, N.: Universality of phenotypic distributions in bacteria. Physical Review Research 6(2), L022043 (2024)

[3] Brenner, N., Braun, E., Yoney, A., Susman, L., Rotella, J., Salman, H.: Single-cell protein dynamics reproduce universal fluctuations in cell populations. The European Physical Journal E 38, 1–9 (2015)

[4] Das, A., Awasthi, P., Sen, R., Kong, W.: Trimmed maximum likelihood estimation for robust learning in generalized linear models (2022)

[5] Duck, P., Marshall, T., Watson, E.: First-passage times for the uhlenbeck-ornstein process. Journal of Physics A: Mathematical and General 19(17), 3545 (1986)

[6] Hadi, A.S., Luceño, A.: Maximum trimmed likelihood estimators: a unified approach, examples, and algorithms. Computational Statistics & Data Analysis 25(3), 251–272 (1997)

[7] Hurwicz, L., Koopmans, T.: Statistical inference in dynamic economic models. Chap. Generalization of the Concept of Identification, edited by TC Koopmans 10, 245–257 (1950)

[8] Jiang, Y., Macrina, A., Peters, G.W.: Multiple barrier-crossings of an ornstein-uhlenbeck diffusion in consecutive periods. Stochastic Analysis and Applications 39(4), 569–609 (2021)

[9] Jun, S., Taheri-Araghi, S.: Cell-size maintenance: universal strategy revealed. Trends in microbiology 23(1), 4–6 (2015)

[10] Leblanc, B., Renault, O., Scaillet, O.: A correction note on the first passage time of an ornstein-uhlenbeck process to a boundary. Finance and Stochastics 4, 109–111 (2000)

[11] Lipton, A., Kaushansky, V.: On the first hitting time density of an ornstein-uhlenbeck process. arXiv preprint arXiv:1810.02390 (2018)

[12] Luo, L., Bai, Y., Fu, X.: Stochastic threshold in cell size control. Physical Review Research 5(1), 013173 (2023)

[13] Neykov, N.M., Müller, C.H.: Breakdown point and computation of trimmed likelihood estimators in generalized linear models. In: Developments in Robust Statistics: International Conference on Robust Statistics 2001. pp. 277–286. Springer (2003)

[14] Peyré, G., Cuturi, M., et al.: Computational optimal transport: With applications to data science. Foundations and Trends® in Machine Learning 11(5-6), 355–607 (2019)

[15] Ricciardi, L.M., Sato, S.: First-passage-time density and moments of the ornstein-uhlenbeck process. Journal of Applied Probability 25(1), 43–57 (1988)

[16] Sassi, A.S., Garcia-Alcala, M., Aldana, M., Tu, Y.: Protein concentration fluctuations in the high expression regime: Taylor’s law and its mechanistic origin. Physical review X 12(1), 011051 (2022)

[17] Shaked, M., Shanthikumar, J.G.: Stochastic orders. Springer (2007)

[18] Si, F., Le Treut, G., Sauls, J.T., Vadia, S., Levin, P.A., Jun, S.: Mechanistic origin of cell-size control and homeostasis in bacteria. Current Biology 29(11), 1760–1770 (2019)

[19] Susman, L., Kohram, M., Vashistha, H., Nechleba, J.T., Salman, H., Brenner, N.: Individuality and slow dynamics in bacterial growth homeostasis. Proceedings of the National Academy of Sciences 115(25), E5679–E5687 (2018)

[20] Taheri-Araghi, S., Bradde, S., Sauls, J.T., Hill, N.S., Levin, P.A., Paulsson, J., Vergassola, M., Jun, S.: Cell-size control and homeostasis in bacteria. Current biology 25(3), 385–391 (2015)

[21] Tanouchi, Y., Pai, A., Park, H., Huang, S., Buchler, N.E., You, L.: Long-term growth data of escherichia coli at a single-cell level. Scientific data 4(1), 170036 (2017)

[22] Tanouchi, Y., Pai, A., Park, H., Huang, S., Stamatov, R., Buchler, N.E., You, L.: A noisy linear map underlies oscillations in cell size and gene expression in bacteria. Nature 523(7560), 357–360 (2015)

[23] Tiruvadi-Krishnan, S., Männik, J., Kar, P., Lin, J., Amir, A., Männik, J.: Coupling between DNA replication, segregation, and the onset of constriction in Escherichia coli. Cell reports 38(12), 110539 (2022)

[24] Vuong, Q.H.: Likelihood ratio tests for model selection and non-nested hypotheses. Econometrica: journal of the Econometric Society pp. 307–333 (1989)

[25] Wang, P., Robert, L., Pelletier, J., Dang, W.L., Taddei, F., Wright, A., Jun, S.: Robust growth of escherichia coli. Current biology 20(12), 1099–1103 (2010)

[26] Wu, F., Dekker, C.: Nanofabricated structures and microfluidic devices for bacteria: from techniques to biology. Chemical Society Reviews 45(2), 268–280 (2016)

[27] Yang, D., Jennings, A.D., Borrego, E., Retterer, S.T., Männik, J.: Analysis of factors limiting bacterial growth in PDMS mother machine devices. Frontiers in microbiology 9, 871 (2018)

[28] Yi, C.: On the first passage time distribution of an ornstein–uhlenbeck process. Quantitative Finance 10(9), 957–960 (2010)

